# Insilico Analysis Reveal Three novel nsSNPs May effect on *GM2A* protein Leading to AB variant of GM2 gangliosidosis

**DOI:** 10.1101/853085

**Authors:** Tebyan A. Abdelhameed, Mujahed I. Mustafa, Dina N. Abdelrahman, Fatima A. Abdelrhman, Mohamed A. Hassan

## Abstract

**Background:** AB variant of GM2 gangliosidosis caused as a result of mutations in GM2 activator gene (*GM2A*) which is regarded as neurodegenerative disorder. This study aimed to predict the possible damaging SNPs of this gene and their impact on the protein using different bioinformatics tools.

**Methods:** SNPs retrieved from the NCBI database were analyzed using several bioinformatics tools. The different tools collectively predicted the effect of single nucleotide substitution on both structure and function of GM2 activator.

**Results:** Three novel mutations were found to be highly damaging to the structure and function of the *GM2A* gene.

**Conclusion:** Four SNPs were found to be highly damaging SNPs that affect function, structure and stability of *GM2A* protein, which they are: C99Y, C112F, C138S and C138R, three of them are novel SNPs (C99Y, C112F and C138S). Also 46 SNPs were predicted to affect miRNAs binding sites on 3’UTR leading to abnormal expression of the resulting protein. These SNPs should be considered as important candidates in causing AB variant of GM2 gangliosidosis and may help in diagnosis and genetic screening of the disease.

## 1. INTRODUCTION

GM2-gangliosidosis is a group of neurological disorders resulting from genetically defective catabolism, and consequent abnormal accumulation, of GM2-ganglioside. Three major types are distinguished: the B variant (Tay-Sachs disease), the O variant (Sandhoff disease), and the AB variant, caused by genetic abnormalities in the genes coding for the beta-hexosaminidase alpha- or beta-subunit, or the GM2-activator protein, respectively. In this study we will focus only in the AB variant of GM2 gangliosidosis.(1, 2) Variant AB of infantile GM2 gangliosidosis is a fatal disease leading invariably to death within the first few years of life, due to the excessive storage of the glycolipids GM2 and GA2 which occurs in the nervous tissue of the patient.(3, 4) The variant AB is characterized by a normal or close to the normal range of this enzyme. Thus, variant AB is an example of a fatal lipid storage disease that is caused not by a defect of a degrading enzyme but rather by a defective factor necessary for the interaction of lipid substrates and the water-soluble hydrolase.(3, 5–7)

The most reported gene for this disorder is *GM2A* (8, 9) which acts as a cofactor for the enzyme. (10, 11) located in chromosome 5q31.3-q33.1 (12–14) The GM2 activator deficiency is caused by mutations in the *GM2A* gene encoding the GM2 activator protein.(15) Different mutations have been reported related to *GM2A,* (16–22) but interestingly, some study shows that, mutation in *GM2A* Leads to a Progressive Chorea-dementia Syndrome.(23)

ELISA system can be used as diagnosis tool as well as therapeutic evaluation of GM2 gangliosidoses using anti-GM2 ganglioside antibodies(24) Can also be diagnosed by demonstrating accumulation of GM2 in the CSF of patients with normal hexosaminidase activity.(25) Biochemical and molecular analysis can also be additional diagnosis approaches.(10, 26–28) Some study revealed that, not all GM2-gangliosidosis pateints can have mutations in the protein coding region of the GM2 activator gene.(29)

The usage of in silico studies has strong impact on the identifcation of candidate SNPs since they are easy and less costly and can facilitate future genetic studies.(30) The main objective of this study was to identify the pathogenic SNPs in coding region and 3′UTR in *GM2A* gene using in silico analysis sofwares and to determine the effect of these SNPs on the structural, functional levels, and regulation of their respective proteins.

## 2. MATERIALS AND METHODS

### 2.1 Data mining

The data on human *GM2A* gene was collected from National Center for Biological Information (NCBI) web site (31). The SNP information (protein accession number and SNP ID) of the MEFV gene was retrieved from the NCBI dbSNP (http://www.ncbi.nlm.nih.gov/snp/) and the protein sequence was collected from Swiss Prot databases (http://expasy.org/)(32).

### 2.2 SIFT

SIFT is a sequence homology-based tool (33) that sorts intolerant from tolerant amino acid substitutions and predicts whether an aminoacid substitution in a protein will have a phenotypic Effect. Considers the position at which the change occurred and the type of amino acid change. Given a protein sequence, SIFT chooses related proteins and obtains an alignment of these proteins with the query. Based on the amino acids appearing at each position in the alignment, SIFT calculates the probability that an amino acid at a position is tolerated conditional on the most frequent amino acid being tolerated. If this normalized value is less than a cutoff, the substitution is predicted to be deleterious. SIFT scores <0.05 are predicted by the algorithm to be intolerant or deleterious amino acid substitutions, whereas scores >0.05 are considered tolerant. It is available at (http://sift.bii.a-star.edu.sg/).

### 2.3 Polyphen-2

It is a software tool (34) to predict possible impact of an amino acid substitution on both structure and function of a human protein by analysis of multiple sequence alignment and protein 3D structure, in addition it calculates position-specific independent count scores (PSIC) for each of two variants, and then calculates the PSIC scores difference between two variants. The higher a PSIC score difference, the higher the functional impact a particular amino acid substitution is likely to have. Prediction outcomes could be classified as probably damaging, possibly damaging or benign according to the value of PSIC as it ranges from (0_1); values closer to zero considered benign while values closer to 1 considered probably damaging and also it can be indicated by a vertical black marker inside a color gradient bar, where green is benign and red is damaging. nsSNPs that predicted to be intolerant by Sift has been submitted to Polyphen as protein sequence in FASTA format that obtained from UniproktB /Expasy after submitting the relevant ensemble protein (ESNP) there, and then we entered position of mutation, native amino acid and the new substituent for both structural and functional predictions. PolyPhen version 2.2.2 is available at (http://genetics.bwh.harvard.edu/pph2/index.shtml).

### 2.4 Provean

Provean is a software tool (35) which predicts whether an amino acid substitution or indel has an impact on the biological function of a protein. it is useful for filtering sequence variants to identify nonsynonymous or indel variants that are predicted to be functionally important. It is available at (https://rostlab.org/services/snap2web/).

### 2.5 SNAP2

Functional effects of mutations are predicted with SNAP2 (36). SNAP2 is a trained classifier that is based on a machine learning device called “neural network”. It distinguishes between effect and neutral variants/non-synonymous SNPs by taking a variety of sequence and variant features into account. The most important input signal for the prediction is the evolutionary information taken from an automatically generated multiple sequence alignment. Also structural features such as predicted secondary structure and solvent accessibility are considered. If available also annotation (i.e. known functional residues, pattern, regions) of the sequence or close homologs are pulled in. In a cross-validation over 100,000 experimentally annotated variants, SNAP2 reached sustained two-state accuracy (effect/neutral) of 82% (at an AUC of 0.9). In our hands this constitutes an important and significant improvement over other methods. It is available at (https://rostlab.org/services/snap2web/).

### 2.6 PHD-SNP

An online Support Vector Machine (SVM) based classifier, is optimized to predict if a given single point protein mutation can be classified as disease-related or as a neutral polymorphism, it is available at: (http://snps.biofold.org/phd-snp/phdsnp.html).

### 2.7 SNP& Go

SNPs&GO is an accurate method that, starting from a protein sequence, can predict whether a variation is disease related or not by exploiting the corresponding protein functional annotation. SNPs&GO collects in unique framework information derived from protein sequence, evolutionary information, and function as encoded in the Gene Ontology terms, and outperforms other available predictive methods. (37) It is available at (http://snps.biofold.org/snps-and-go/snps-and-go.html)

### 2.8 P-Mut

PMUT a web-based tool (38) for the annotation of pathological variants on proteins, allows the fast and accurate prediction (approximately 80% success rate in humans) of the pathological character of single point amino acidic mutations based on the use of neural networks. It is available at (http://mmb.irbbarcelona.org/PMut).

### 2.9 I-Mutant 3.0

I-Mutant 3.0 Is a neural network based tool (39) for the routine analysis of protein stability and alterations by taking into account the single-site mutations. The FASTA sequence of protein retrieved from UniProt is used as an input to predict the mutational effect on protein stability. It is available at (http://gpcr2.biocomp.unibo.it/cgi/predictors/I-Mutant3.0/I-Mutant3.0.cgi).

### 2.10 Project Hope

Online software is available at: (http://www.cmbi.ru.nl/hope/method/). It is a web service where the user can submit a sequence and mutation. The software collects structural information from a series of sources, including calculations on the 3D protein structure, sequence annotations in UniProt and prediction from other software. It combines this information to give analysis for the effect of a certain mutation on the protein structure. HOPE will show the effect of that mutation in such a way that even those without a bioinformatics background can understand it. It allows the user to submit a protein sequence (can be FASTA or not) or an accession code of the protein of interest. In the next step, the user can indicate the mutated residue with a simple mouse click. In the final step, the user can simply click on one of the other 19 amino acid types that will become the mutant residue, and then full report well is generated (40).

### 2.11 PolymiRTS

PolymiRTS is a software used to predict 3UTR (un-translated region) polymorphism in microRNAs and their target sites available at (http://compbio.uthsc.edu/miRSNP/). It is a database of naturally occurring DNA variations in mocriRNAs (miRNA) seed region and miRNA target sites. MicroRNAs pair to the transcript of protein coding genes and cause translational repression or mRNA destabilization. SNPs in microRNA and their target sites may affect miRNA-mRNA interaction, causing an effect on miRNA-mediated gene repression, PolymiRTS database was created by scanning 3UTRs of mRNAs in human and mouse for SNPs in miRNA target sites. Then, the effect of polymorphism on gene expression and phenotypes are identified and then linked in the database. The PolymiRTS data base also includes polymorphism in target sites that have been supported by a variety of experimental methods and polymorphism in miRNA seed regions. (42)

### 2.12 UCSF Chimera (University of California at San Francisco)

UCSF Chimera (https://www.cgl.ucsf.edu/chimera/) is a highly extensible program for interactive visualization and analysis of molecular structures and related data, including density maps, supramolecular assemblies, sequence alignments, docking results, trajectories, and conformational ensembles. High-quality images and animations can be generated. Chimera includes complete documentation and several tutorials. Chimera is developed by the Resource for Biocomputing, Visualization, and Informatics (RBVI), supported by the National Institutes of Health (P41-GM103311).(41)

### 2.13 Raptor X

RaptorX (http://raptorx.uchicago.edu/): It is a web server predicting structure property of a protein sequence without using any templates. It outperforms other servers, especially for proteins without close homologs in PDB or with very sparse sequence profile. The server predicts tertiary structure (43)

### 2.14 GeneMANIA

It is gene interaction software that finds other genes which is related to a set of input genes using a very large set of functional association data. Association data include protein and genetic interactions, pathways, co-expression, co-localization and protein domain similarity. GeneMANIA also used to find new members of a pathway or complex, find additional genes you may have missed in your screen or find new genes with a specific function, such as protein kinases. available at (https://genemania.org/) (44).

## 3. RESUTS

### 3.1 Retrieval of SNPs from the Database

All SNPs data related to *GM2A* gene was gathered from dbSNP in National Center of Biotechnology Information (NCBI) database. This gene containing a total of 247 SNPs in coding region, of which 159 were missense, 74 synonymous, 5 nonsense, 8 frame shift, and 676 in 3’untranslated region (3’ UTR).

**Figure (1):**
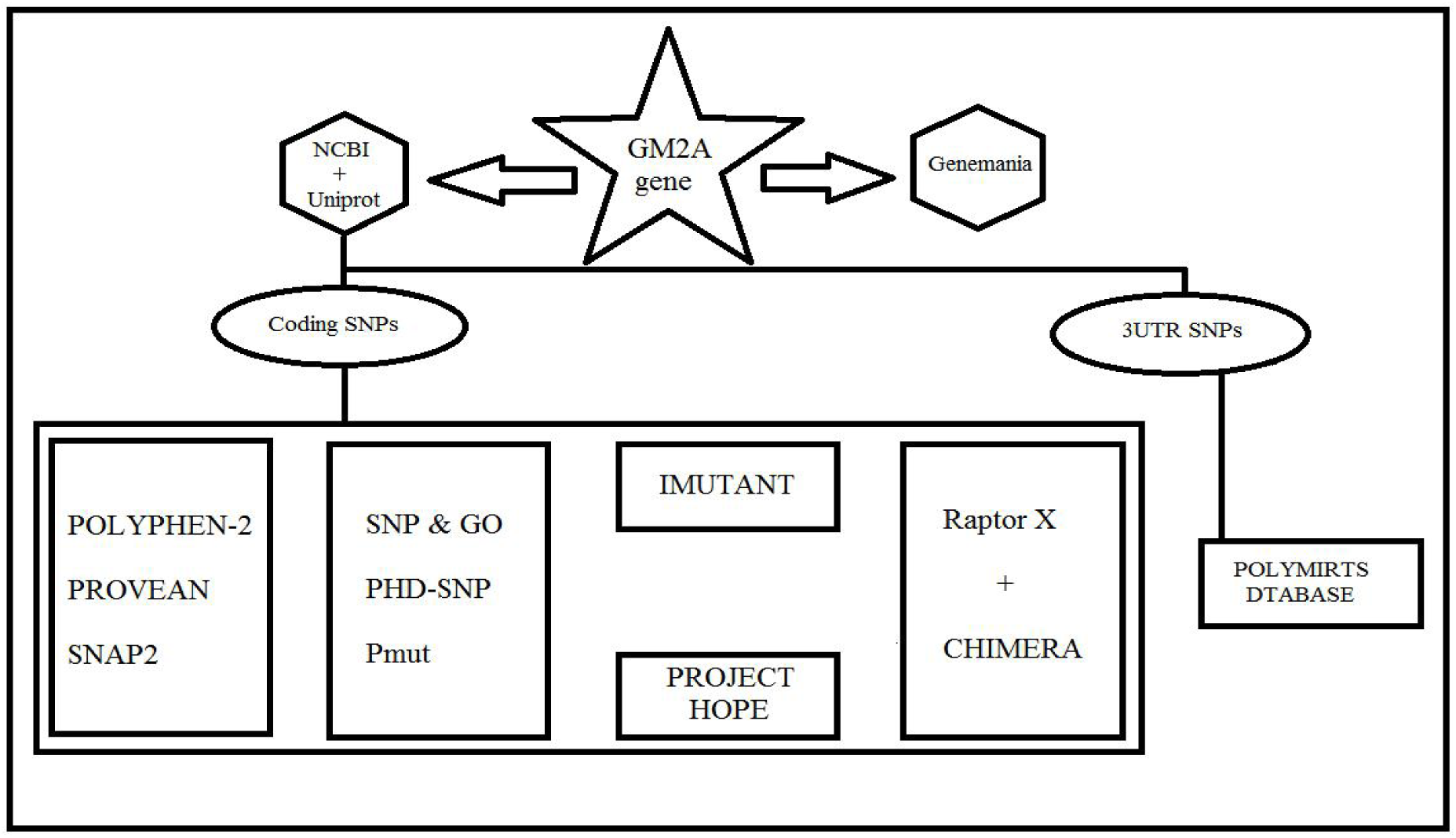
Flowchart of the software used in the analysis processes.

### 3.2 The functional effect of Deleterious and damaging nsSNPs of *GM2A* by SIFT, Provean PolyPhen-2, and SNAP2

**Table (1):** Prediction of functional effect of Deleterious and damaging nsSNPs by SIFT, Polyphen-2, PROVEAN and SNAP2.

| Variant | Polyphen2 Prediction | Polyphen2 score | PROVEAN score | PROVEAN Prediction (cutoff= -2.5) | SNAP2 Prediction | SNAP2 score |
| --- | --- | --- | --- | --- | --- | --- |
| C99Y | possibly damaging | 1 | -10.833 | Deleterious | effect | 85 |
| C112F | probably damaging | 1 | -10.8 | Deleterious | effect | 81 |
| C138R | probably damaging | 1 | -12 | Deleterious | effect | 89 |
| C138S | probably damaging | 1 | -10 | Deleterious | effect | 84 |

### 3.3 Functional analysis of ADAMTS13 gene using Disease Related and pathological effect of nsSNPs by PhD-SNP, SNPs & GO and PMut softwares

**Table (2):** Prediction of disease related nsSNPs through using: PhD-SNP, SNPs & GO and PMut softwares.

| Mutation | SNP Prediction | R I | Probability | PhD Prediction | R I | Probability | Pmut Position | Mutation | Prediction |
| --- | --- | --- | --- | --- | --- | --- | --- | --- | --- |
| C99Y | Disease | 7 | 0.831 | Disease | 9 | 0.949 | 99 | C → Y<br>(Cys → Tyr) | 0.82 (90%)<br>**Disease** |
| C112F | Disease | 3 | 0.67 | Disease | 8 | 0.887 | 112 | C → F<br>(Cys → Phe) | 0.82 (90%)<br>**Disease** |
| C138R | Disease | 3 | 0.67 | Disease | 8 | 0.896 | 138 | C → R<br>(Cys → Arg) | 0.82 (90%)<br>**Disease** |
| C138S | Disease | 2 | 0.623 | Disease | 7 | 0.843 | 138 | C → S<br>(Cys → Ser) | 0.82 (90%)<br>**Disease** |

### 3.4 Prediction of Change in Stability due to Mutation Using I-Mutant 3.0 Server

**Table (3):** Effect of nsSNPs on protein stability using I-Mutant

3.4 Prediction of Change in Stability due to Mutation Using I-Mutant 3.0 Server **Table (3):** Effect of nsSNPs on protein stability using I-Mutant
| dbSNP rs# | Mutation | Prediction | DDG value (score) |
| --- | --- | --- | --- |
| rs751417546 | C99Y | Increase Protein Stability | -0.05 |
| rs773743799 | C112F | Decrease Protein Stability | 0.03 |
| rs137852797 | C138R | Decrease Protein Stability | -0.24 |
| rs1174735558 | C138S | Decrease Protein Stability | -0.82 |

### 3.5 Modeling of amino acid substitution effects on protein structure using Chimera and Project Hope Softwares

**Figure (2):**
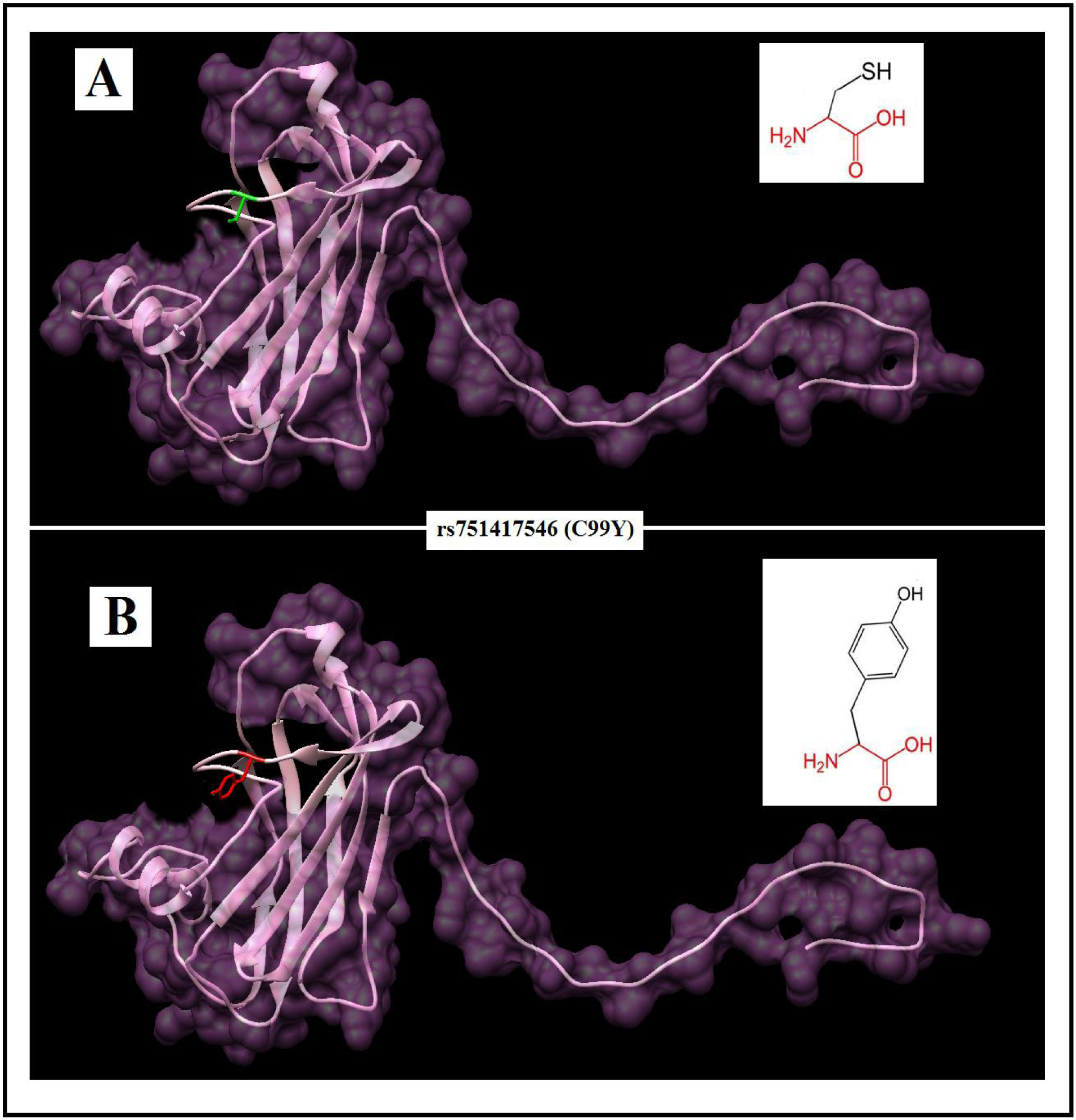
Effect of C99Y (rs751417546) SNP on protein structure in which Cysteine mutated into Tyrosine at position 99 using Project HOPE and Chimera.

**Figure (3):**
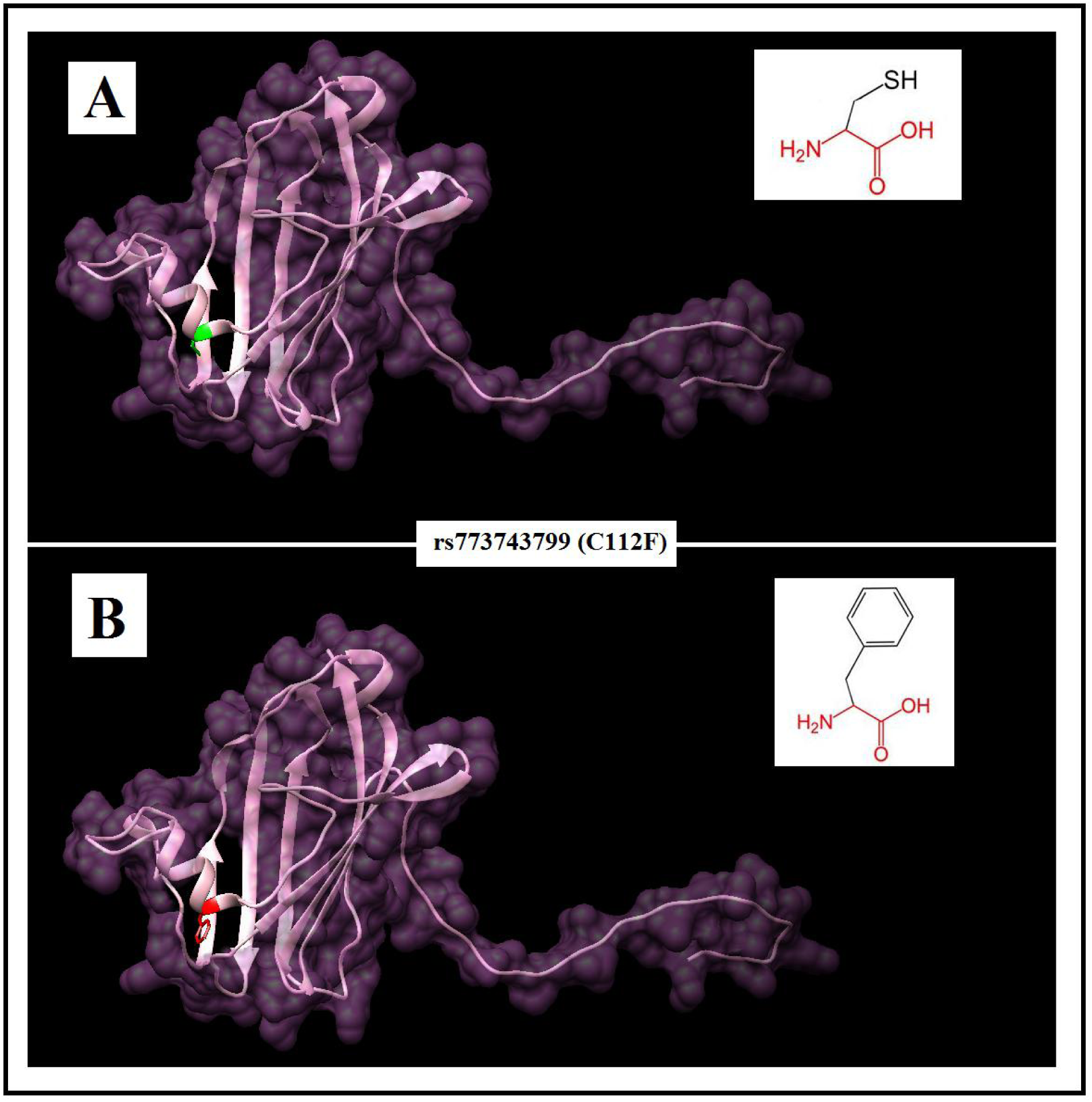
Effect of C112F (rs773743799) SNP on protein structure in which Cysteine mutated into Phenylalanine at position 112 using project HOPE and Chimera.

**Figure (4).**
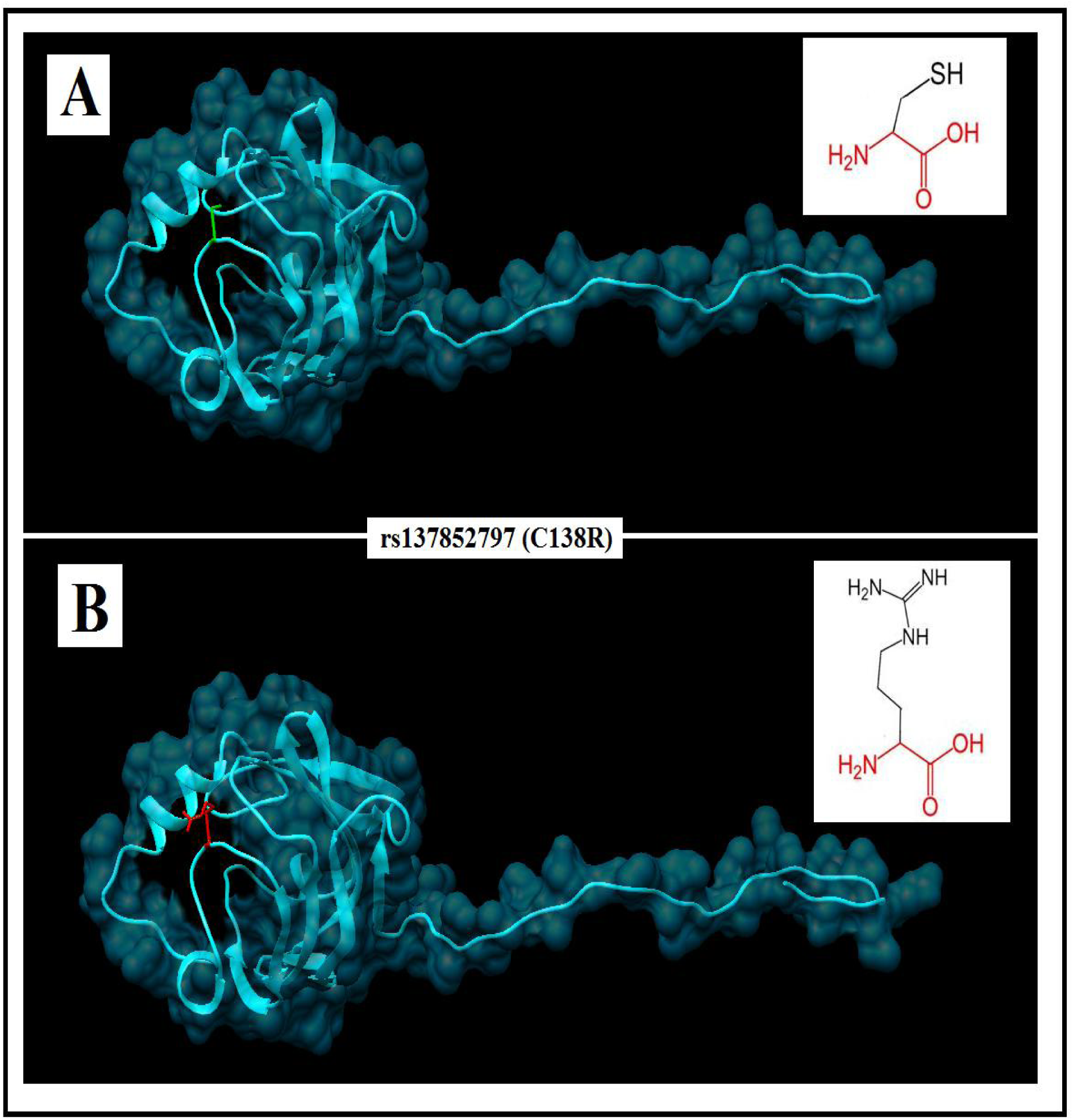
Effect of C138R (rs137852797) SNP on protein structure in which Cysteine mutated into Arginine at position 138 using project HOPE and Chimera.

**Figure (5).**
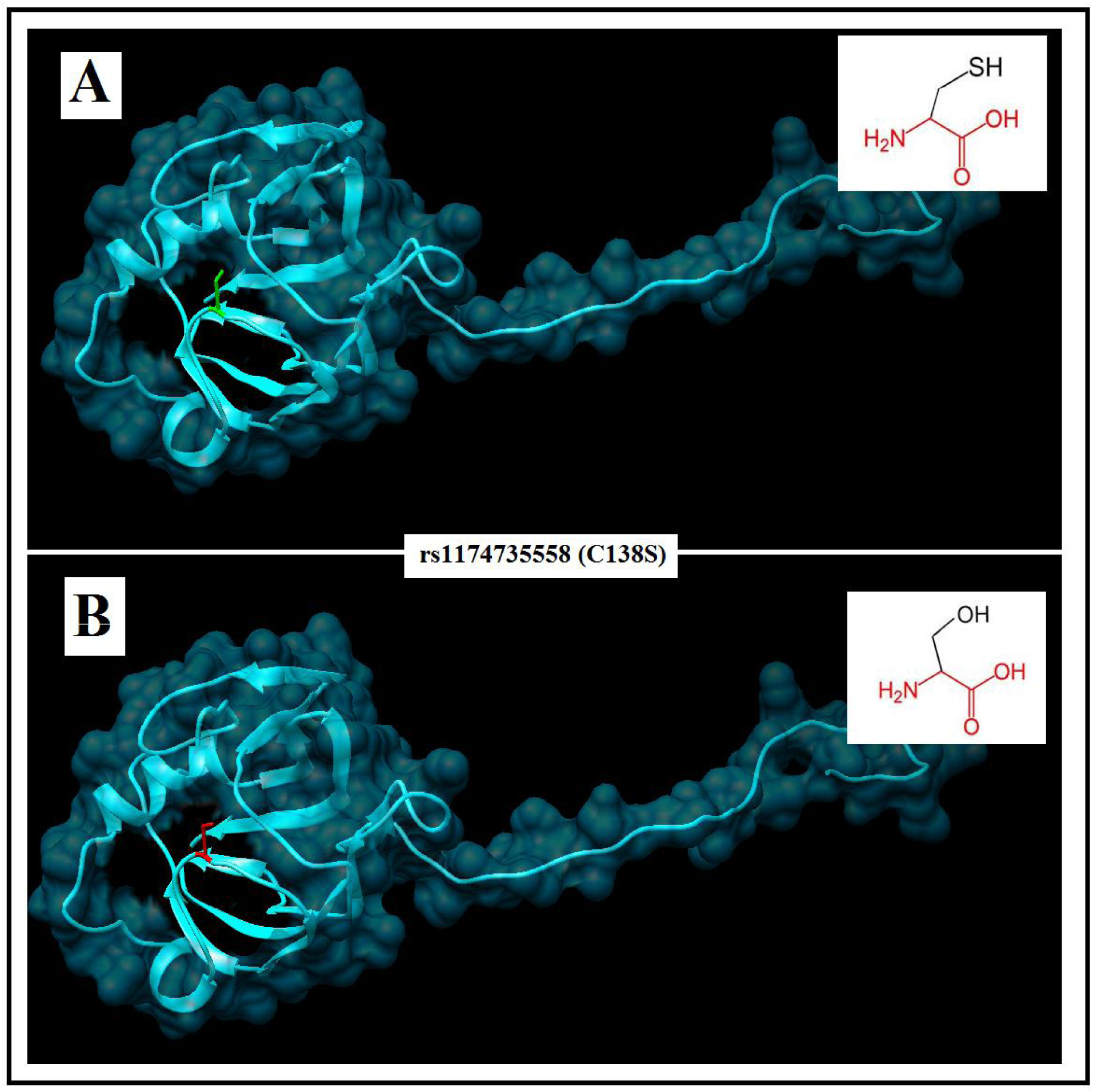
Effect of C138S (rs137852797) SNP on protein structure in which Cysteine mutated into Serine at position 138 using project HOPE and Chimera.

### 3.6 SNPs effect on 3’UTR Region (miRNA binding sites) in *GM2A* using PolymiRTS Database

**Table (4):**
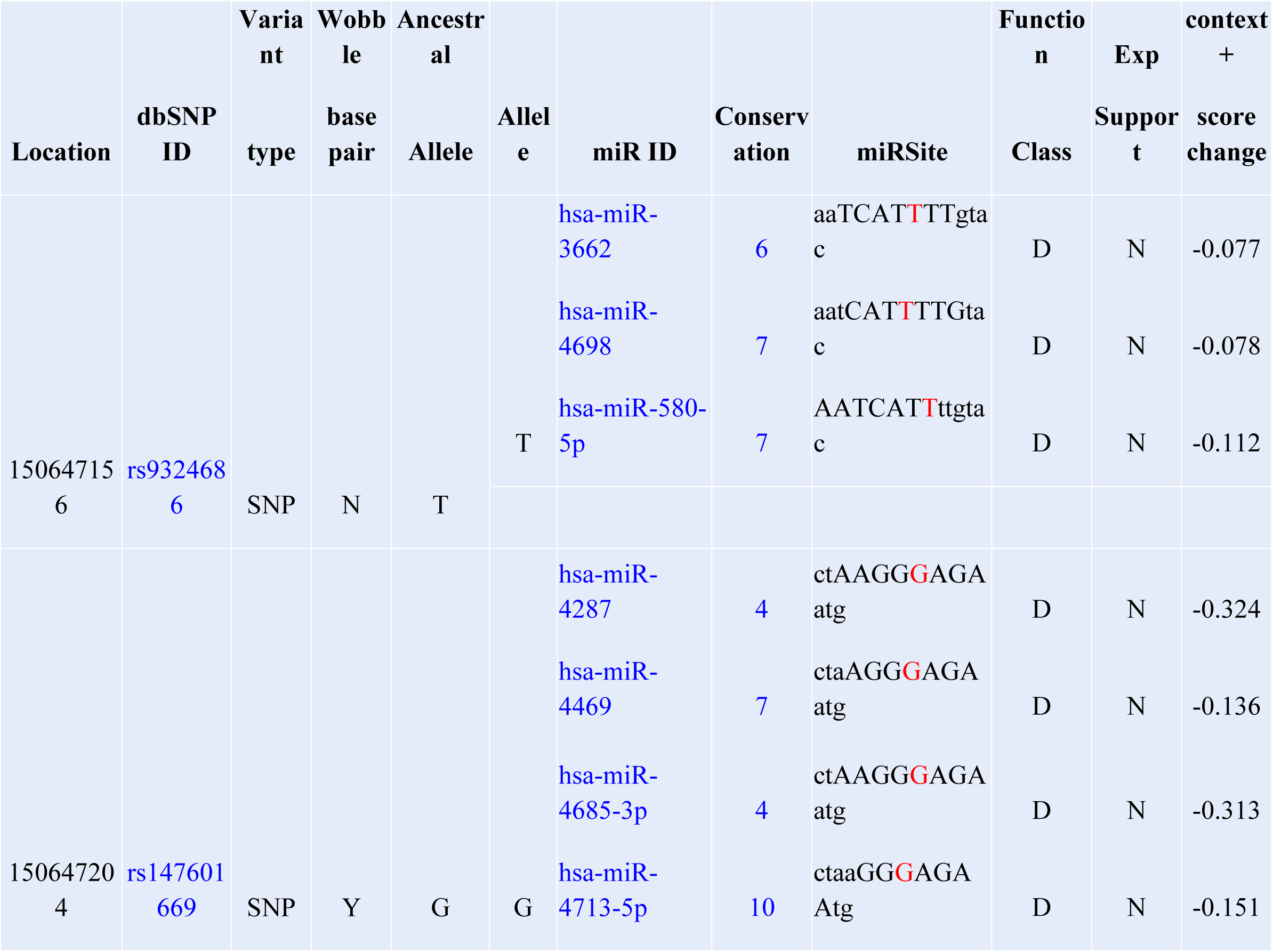
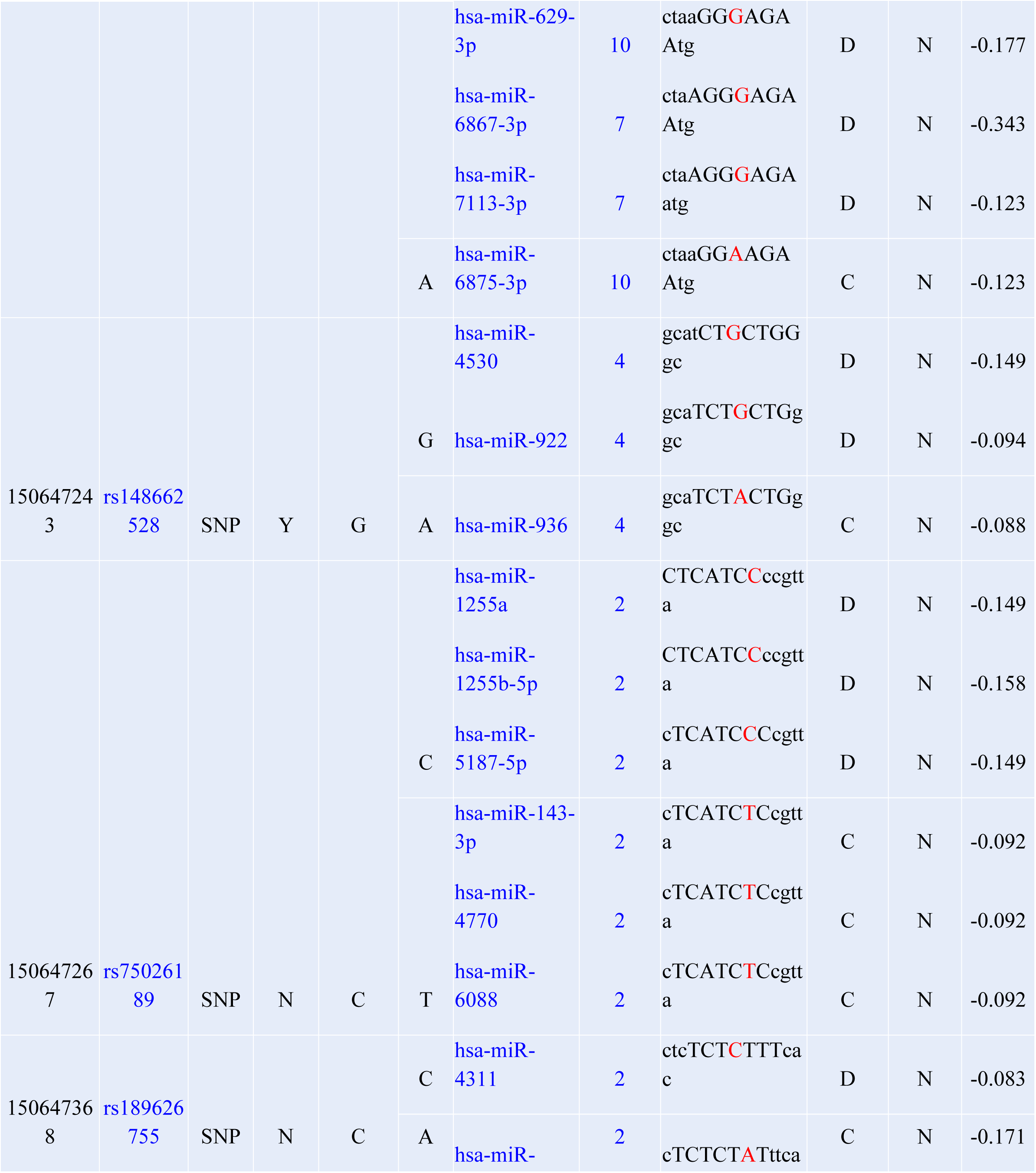
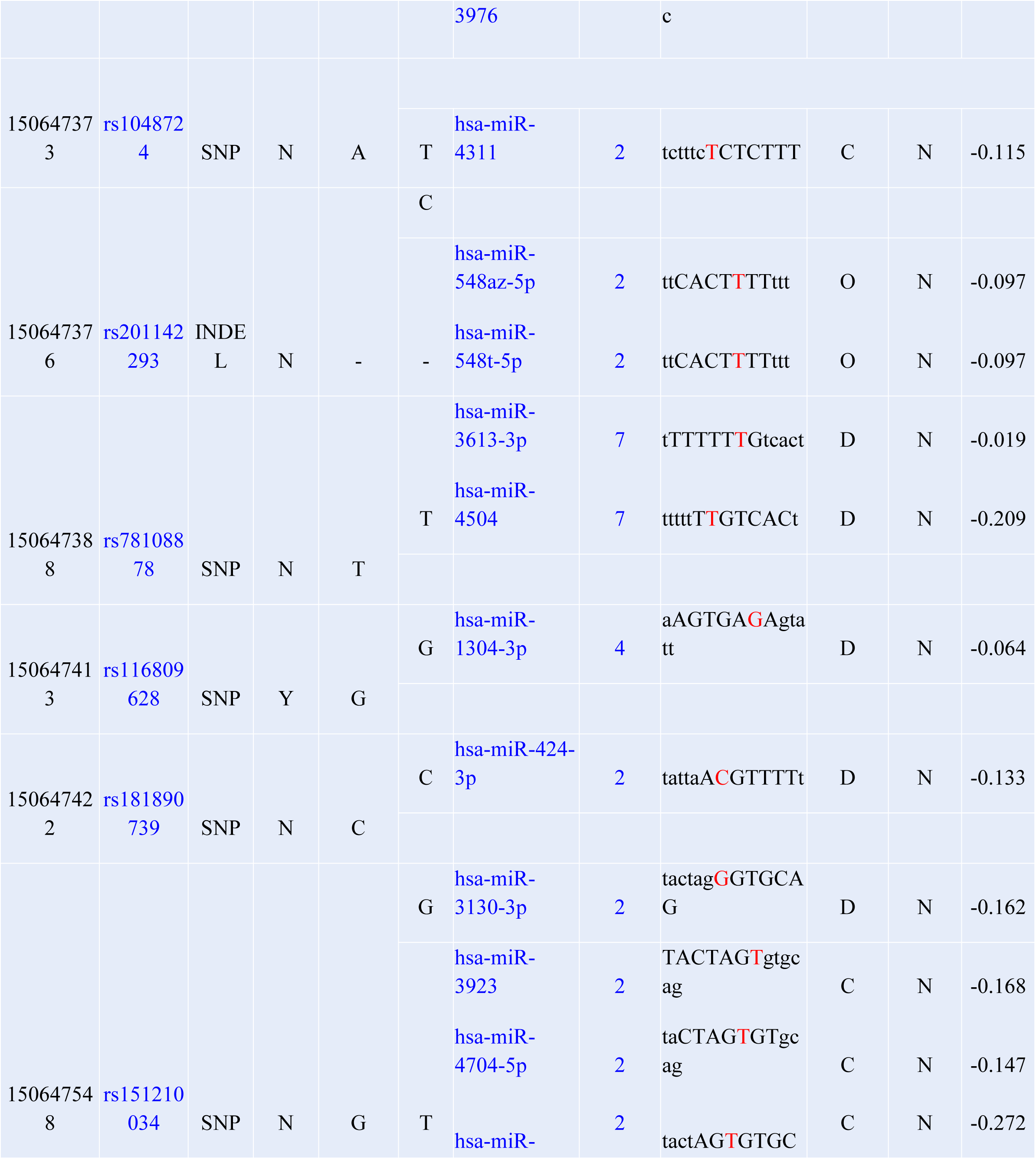
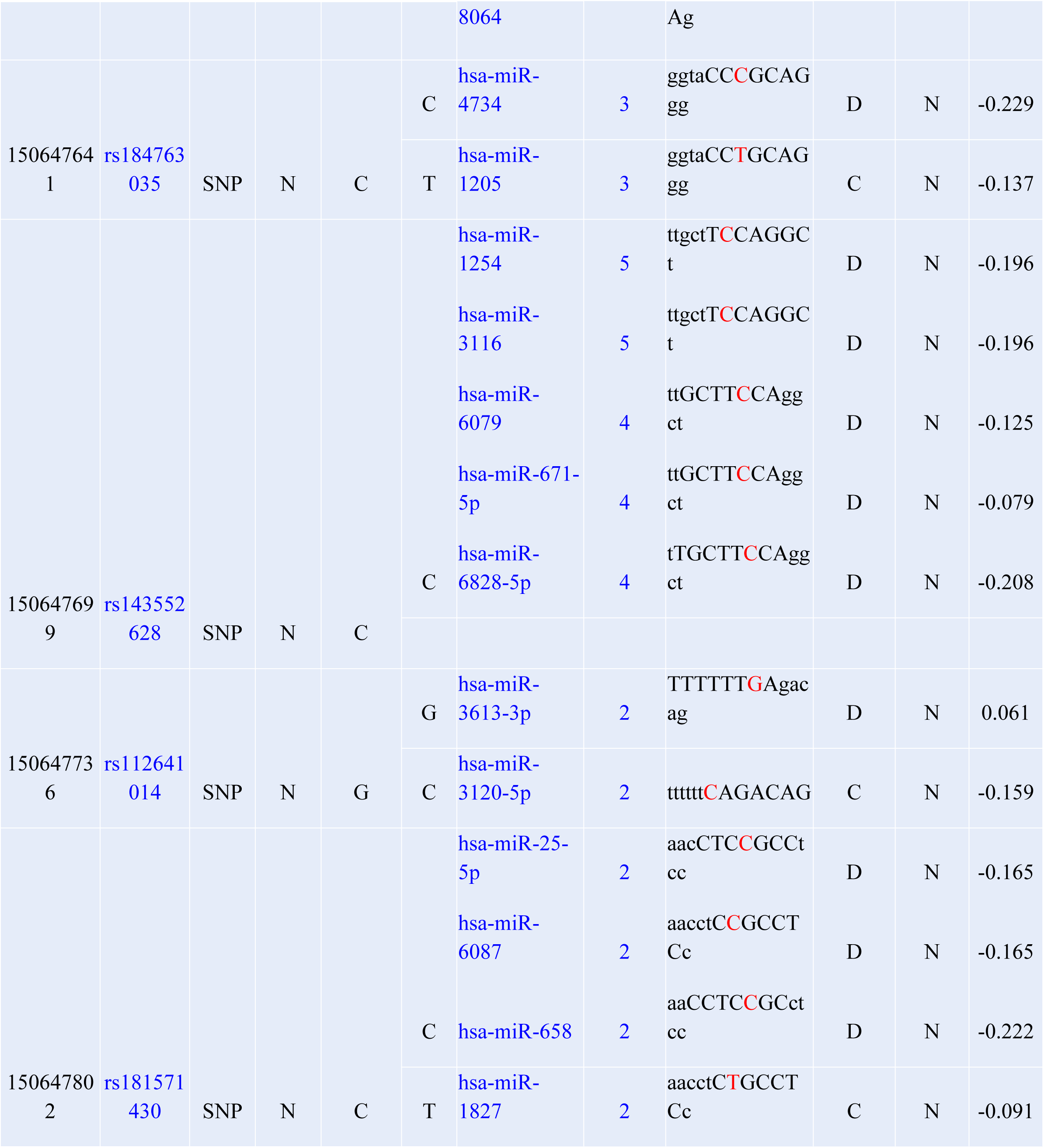
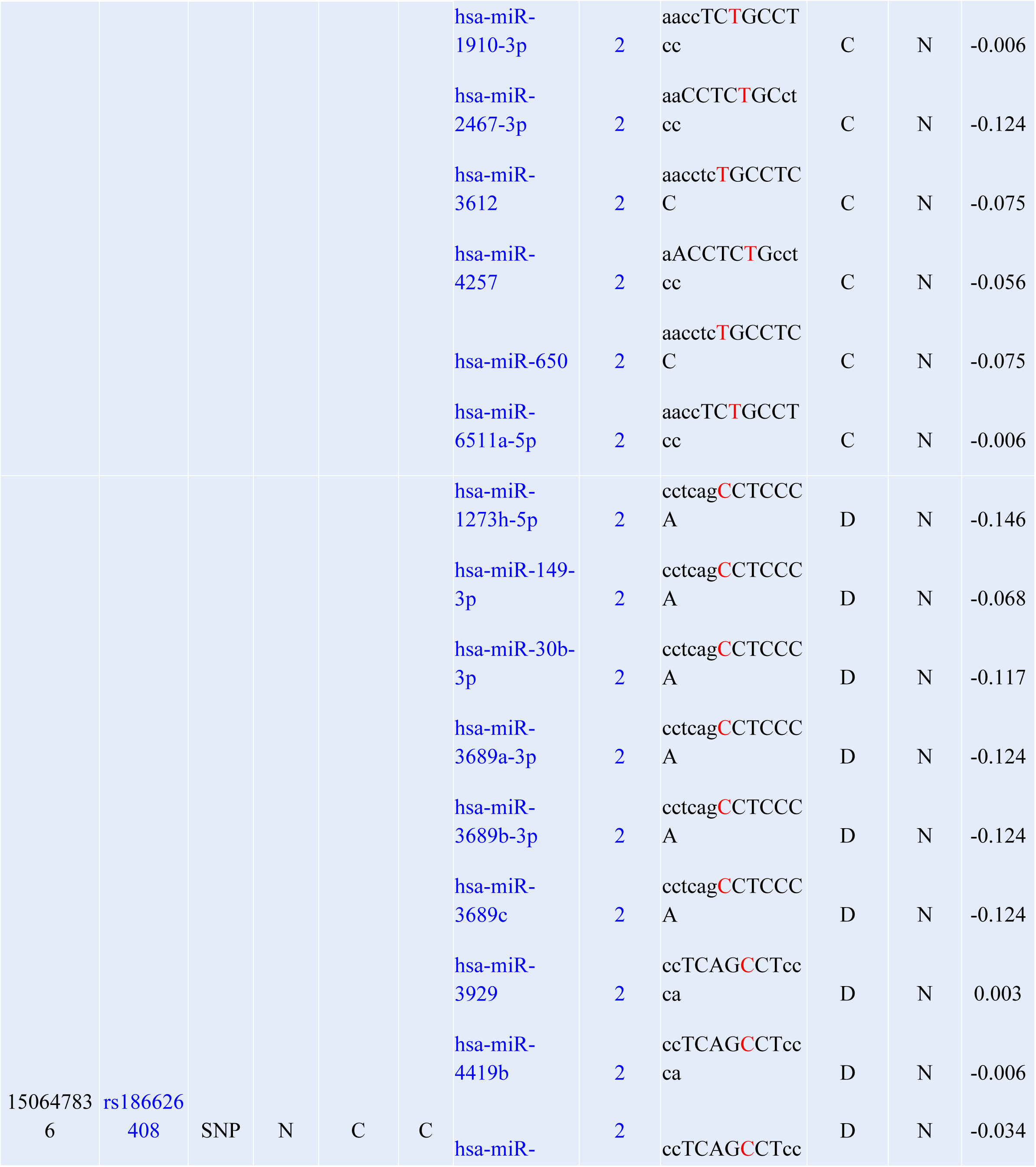
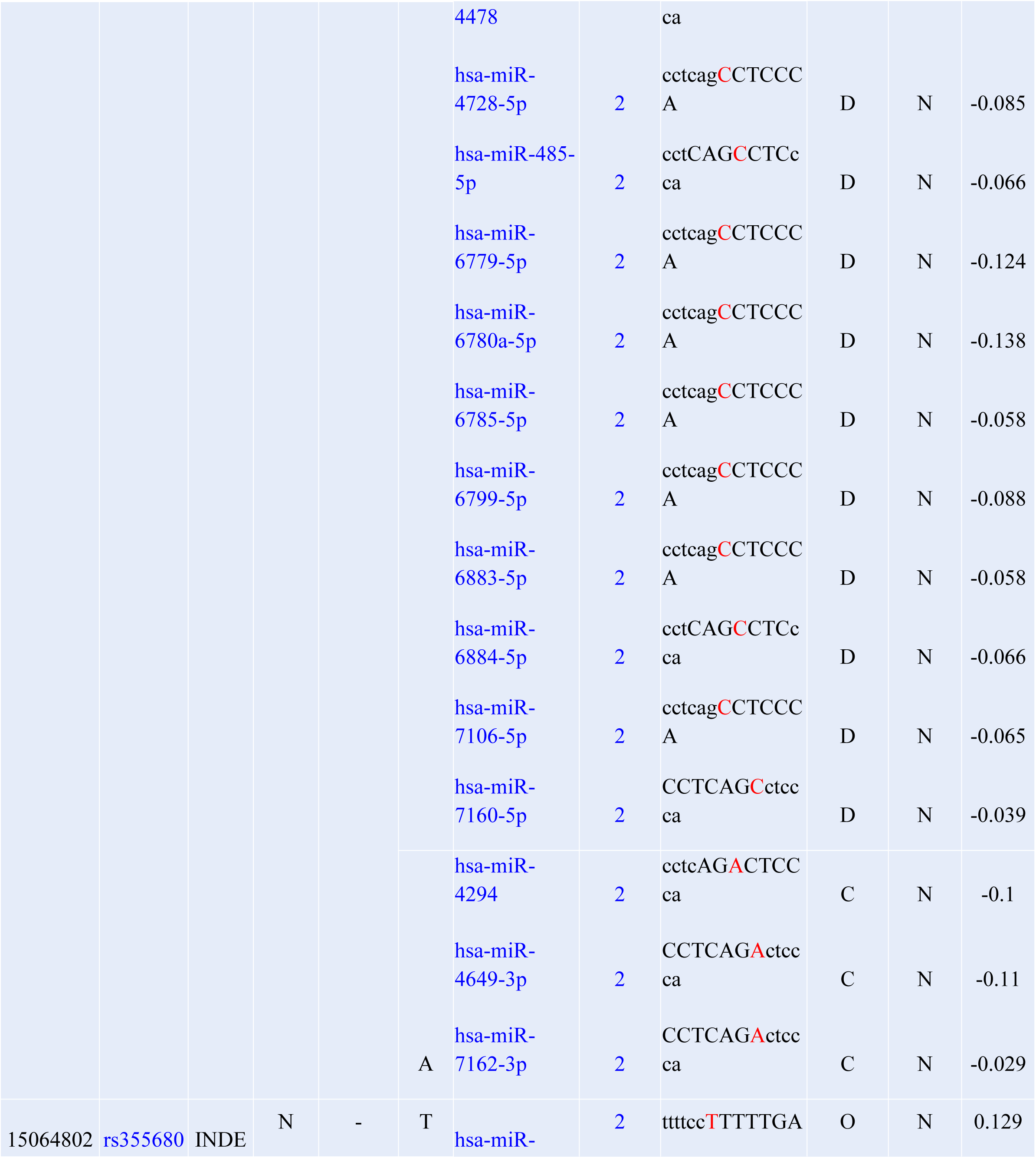
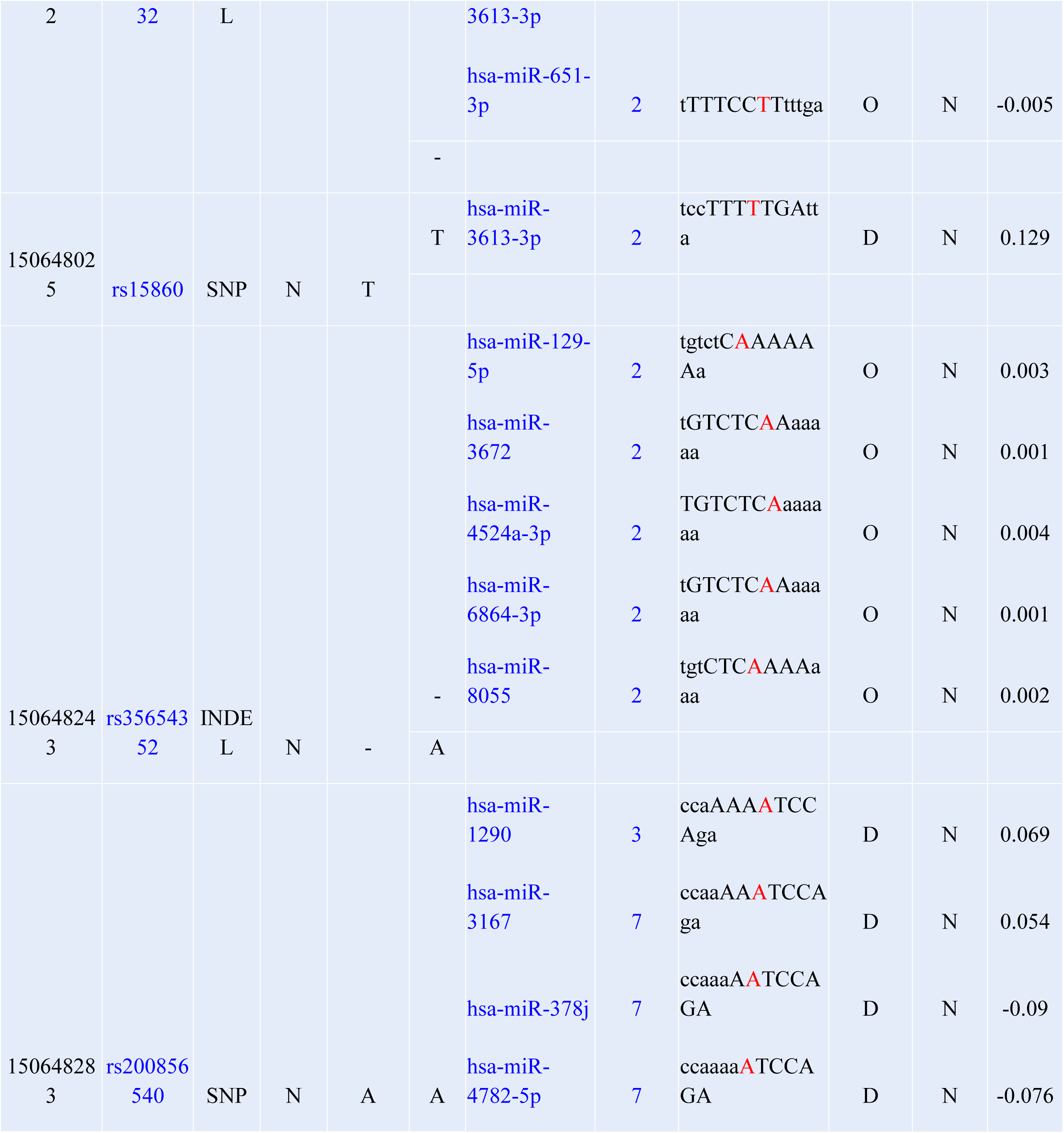
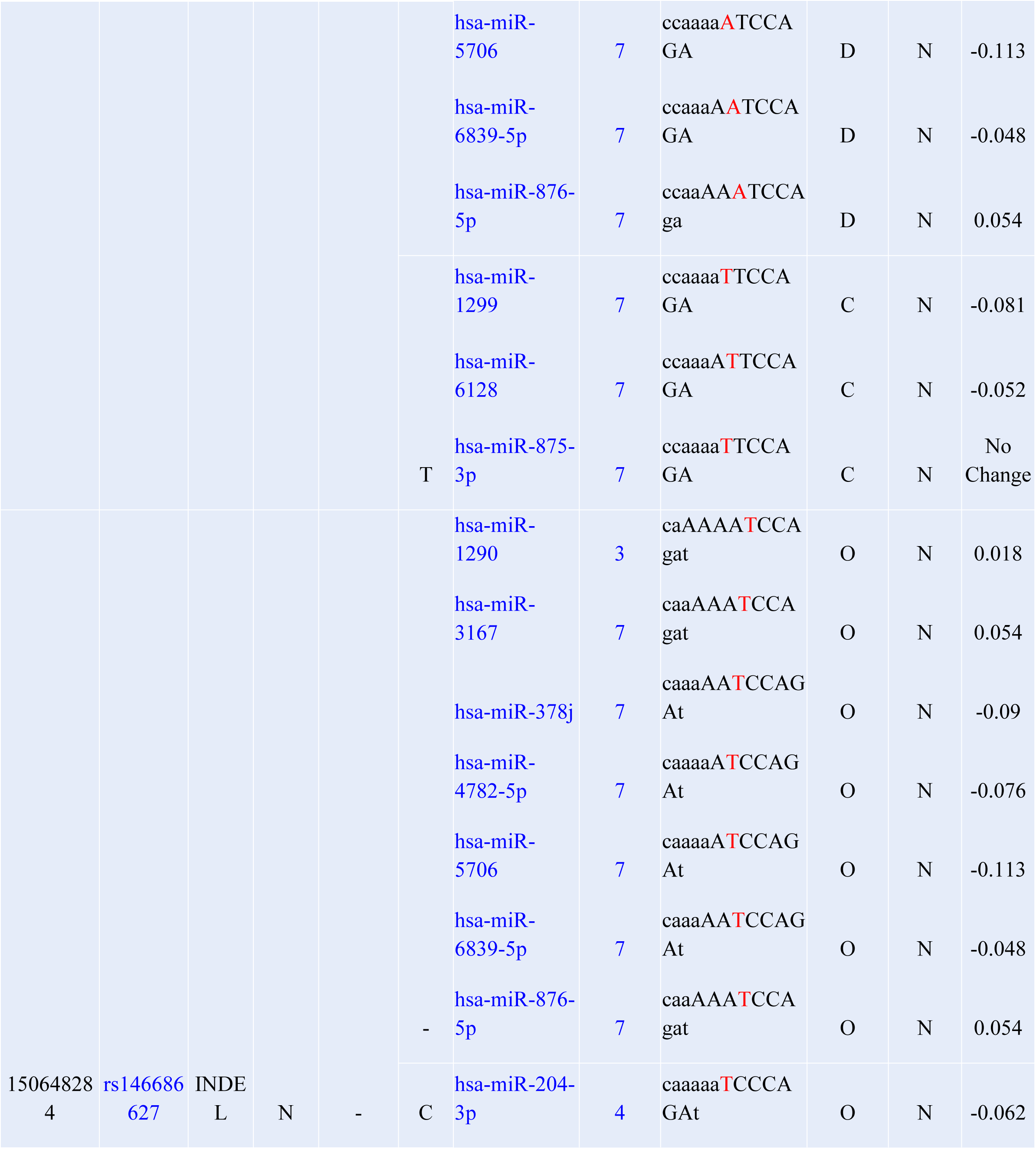
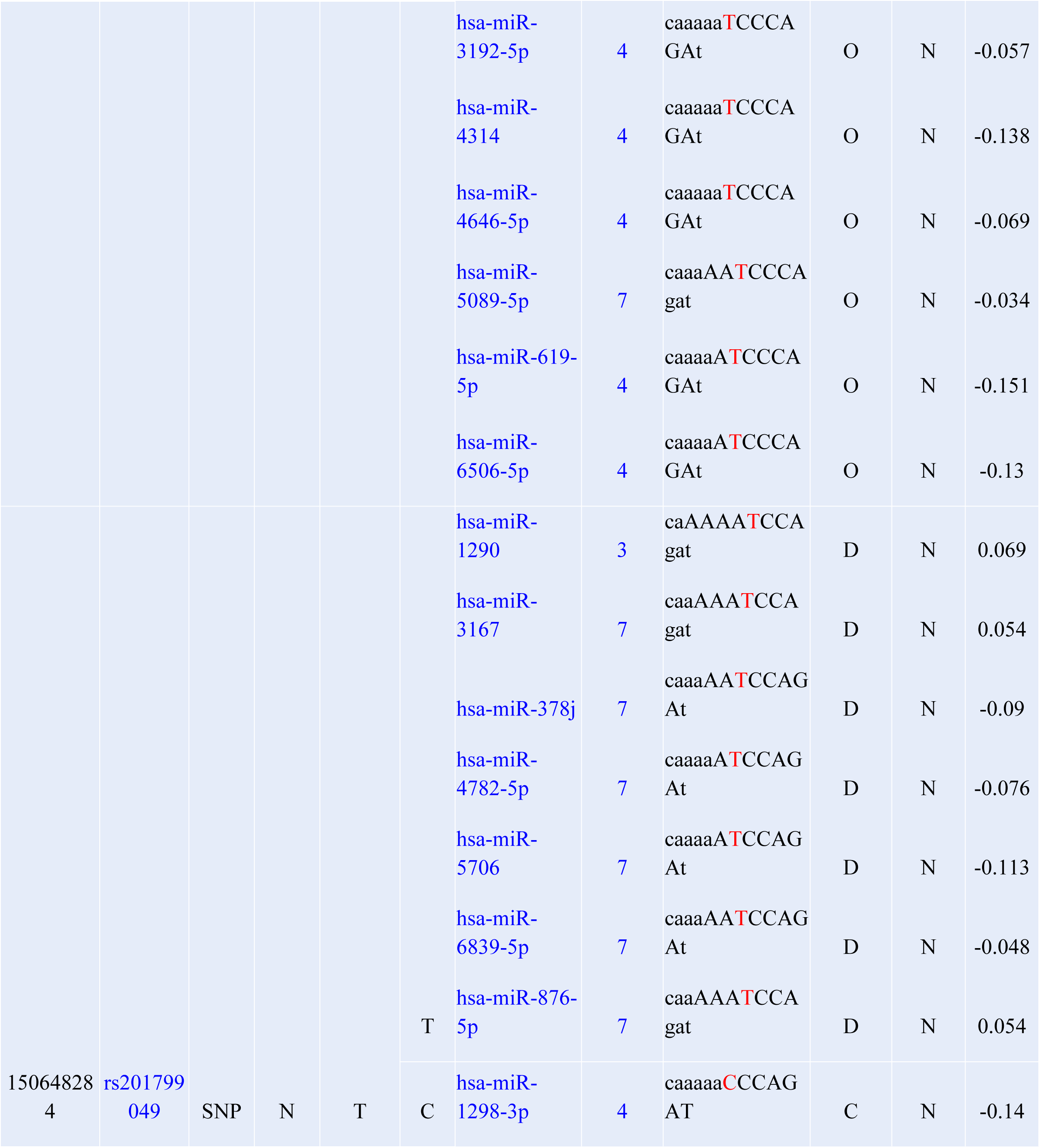
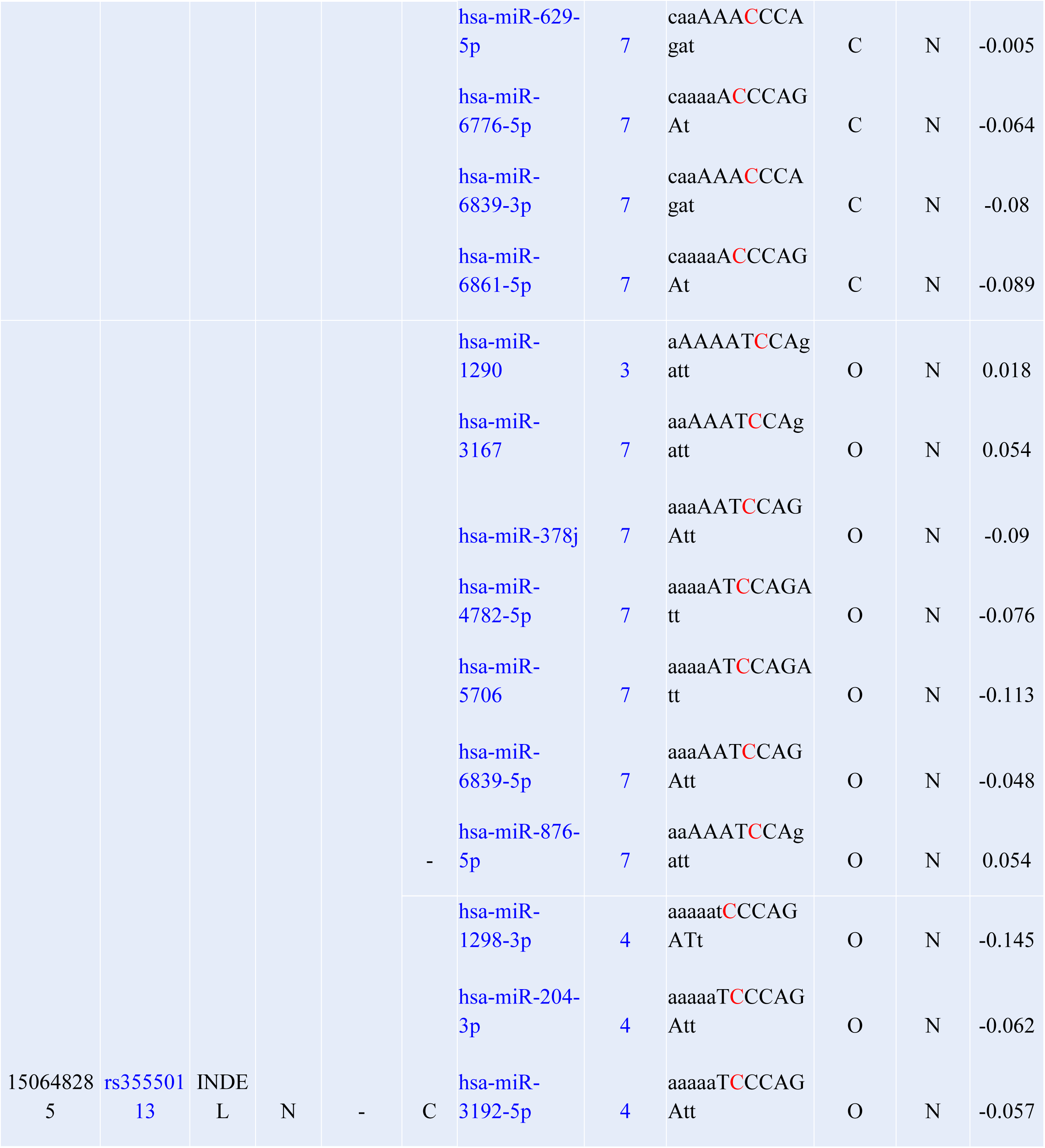
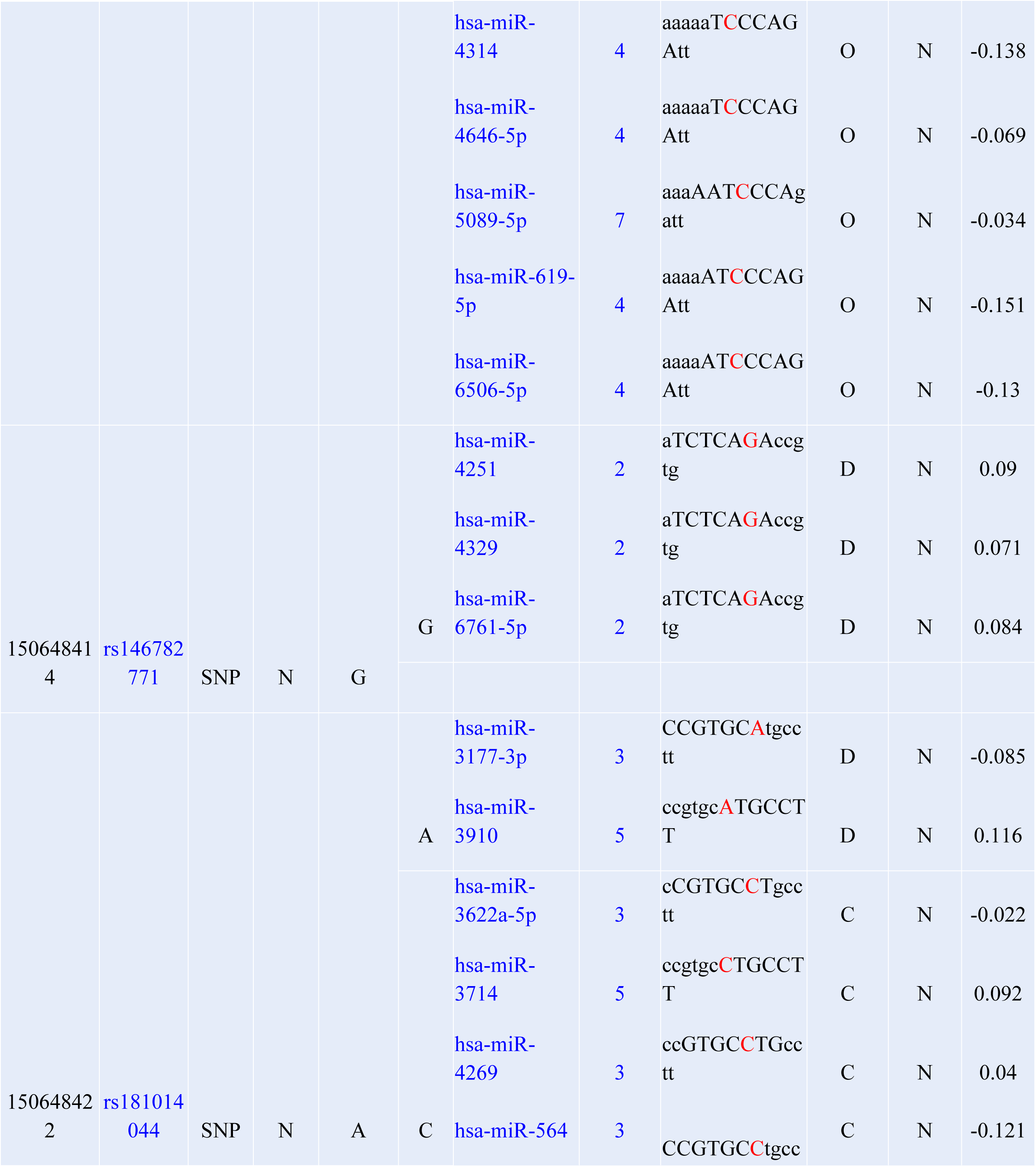
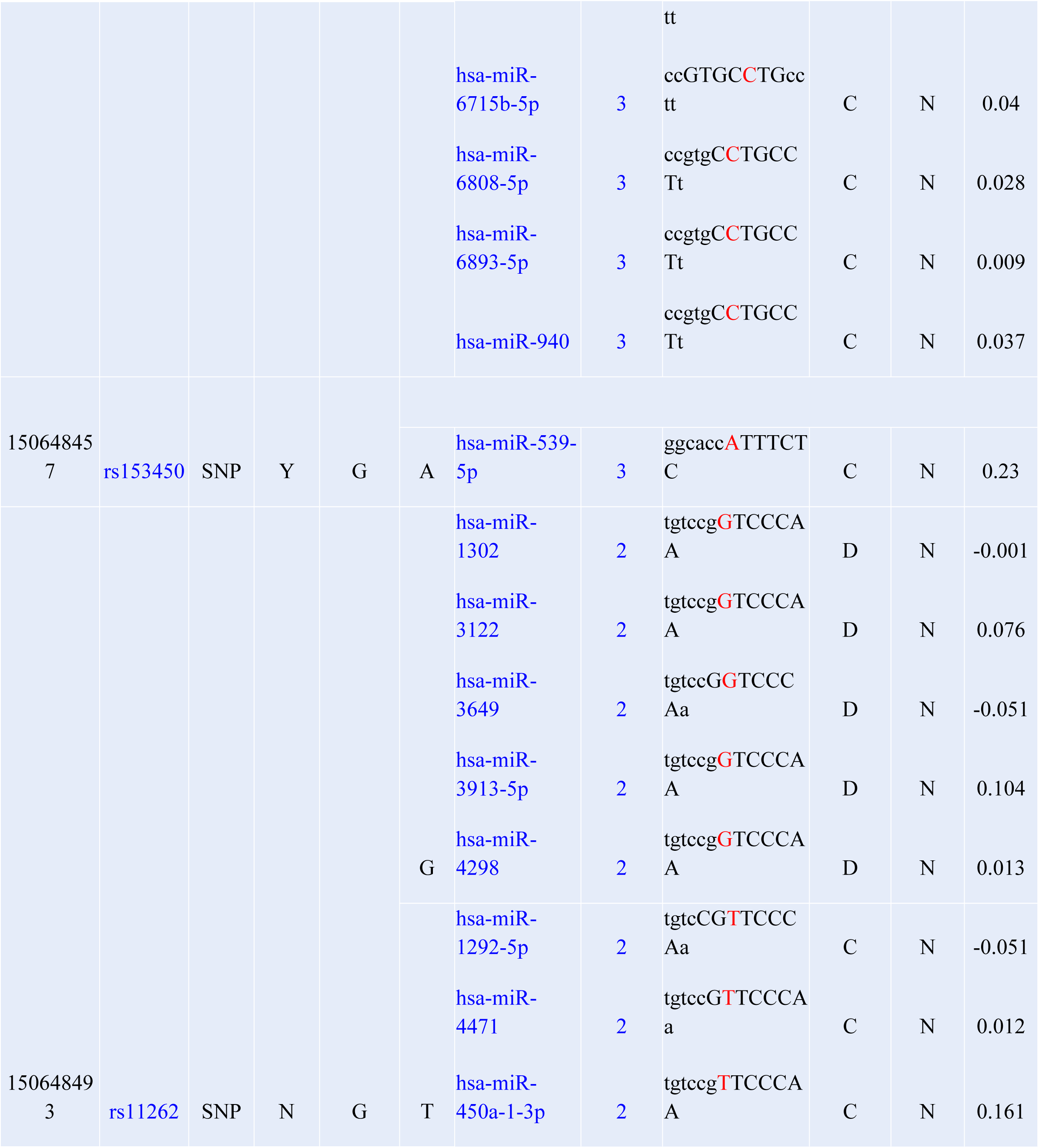
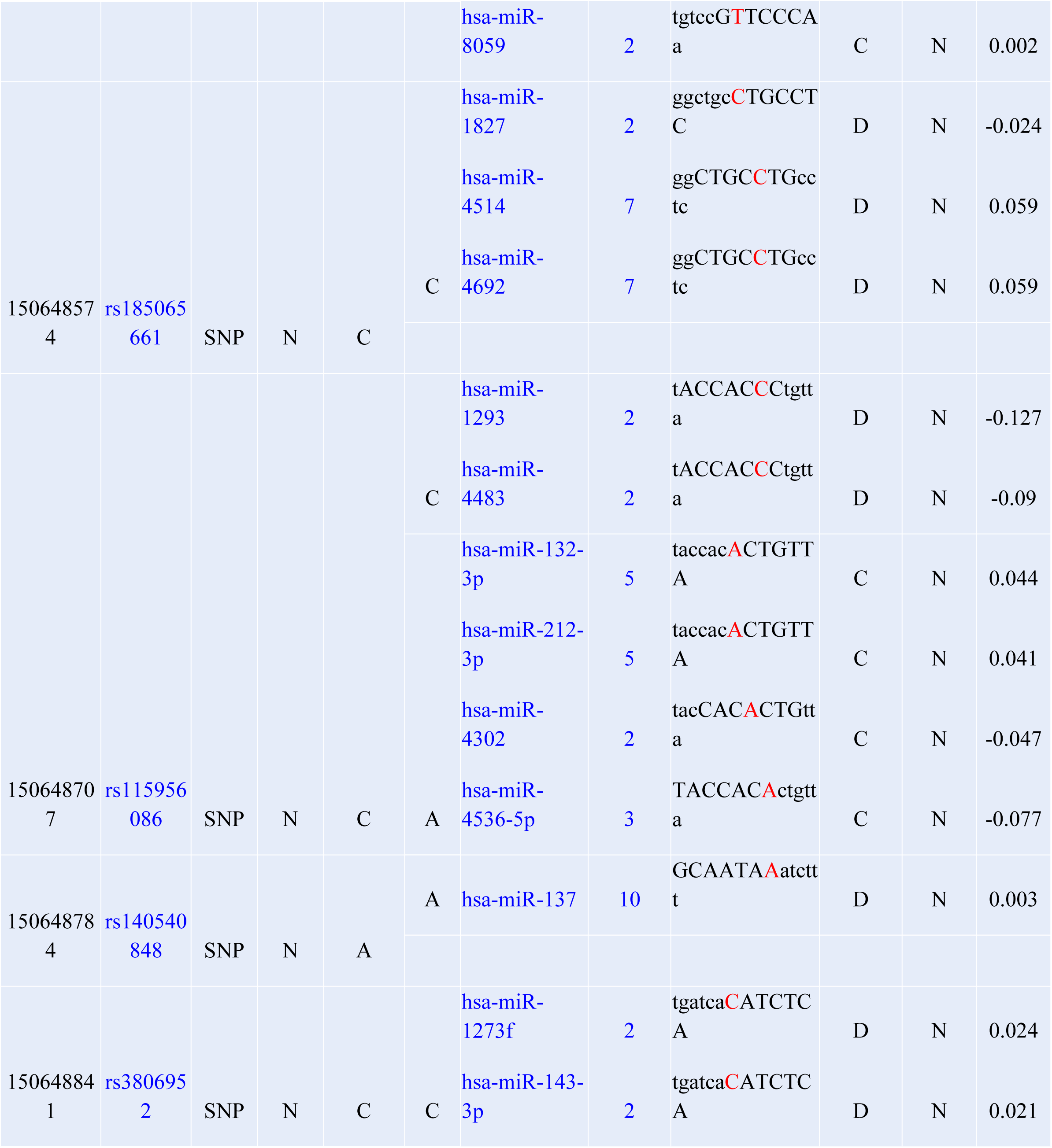
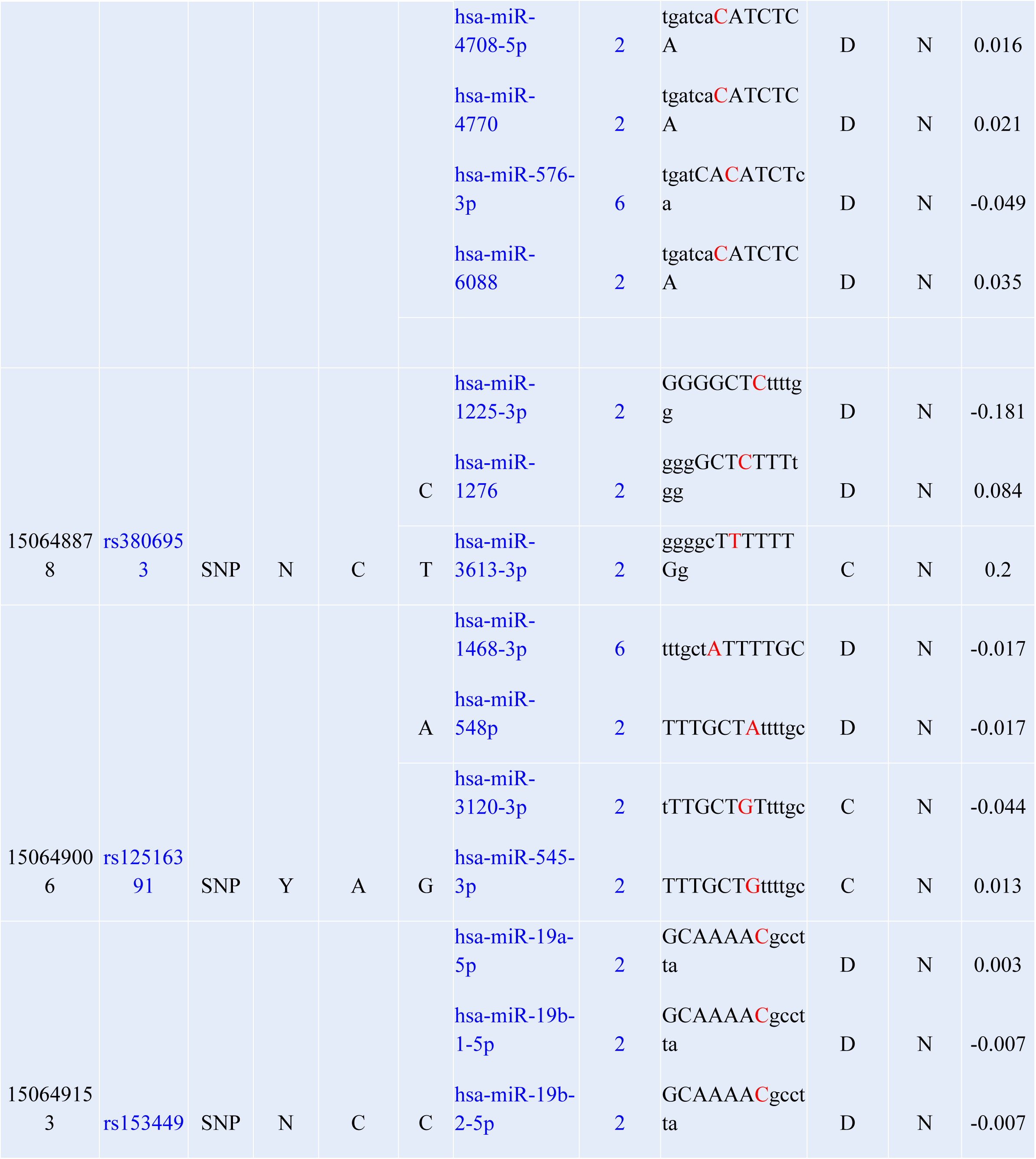
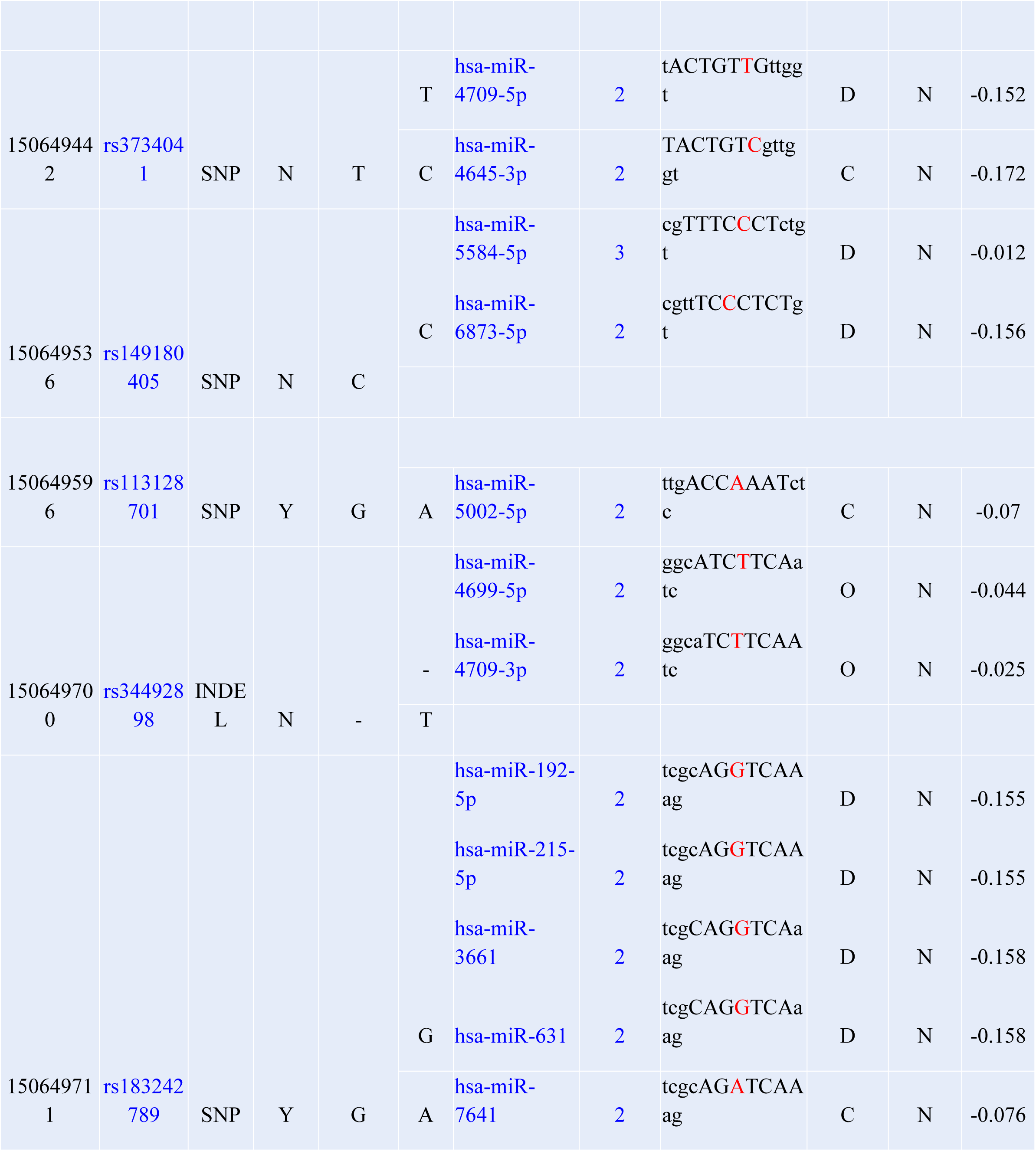
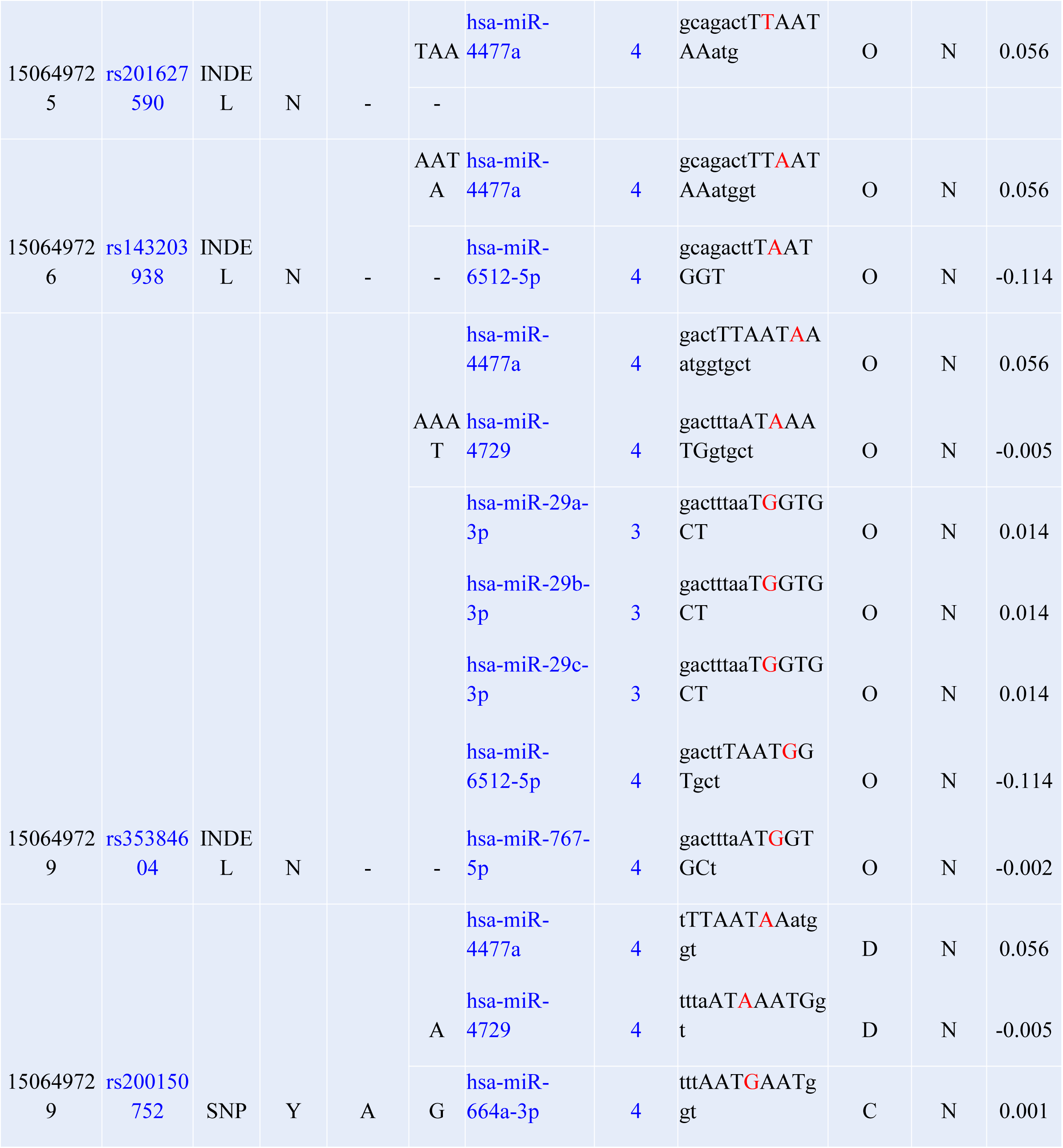
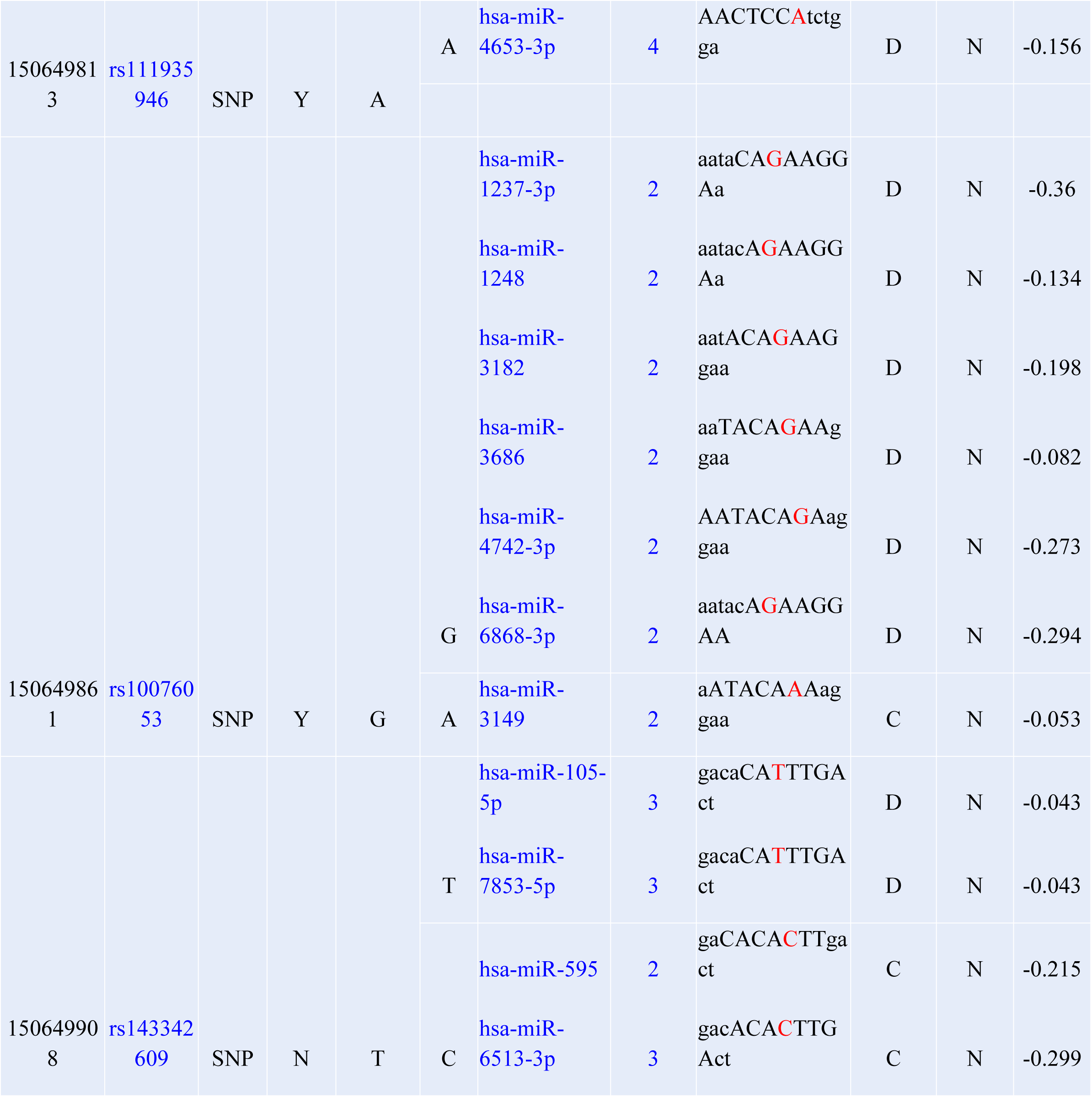
prediction of SNPs sites in *GM2A* gene at the 3’UTR Region using PolymiRTS

**Figure 3:**
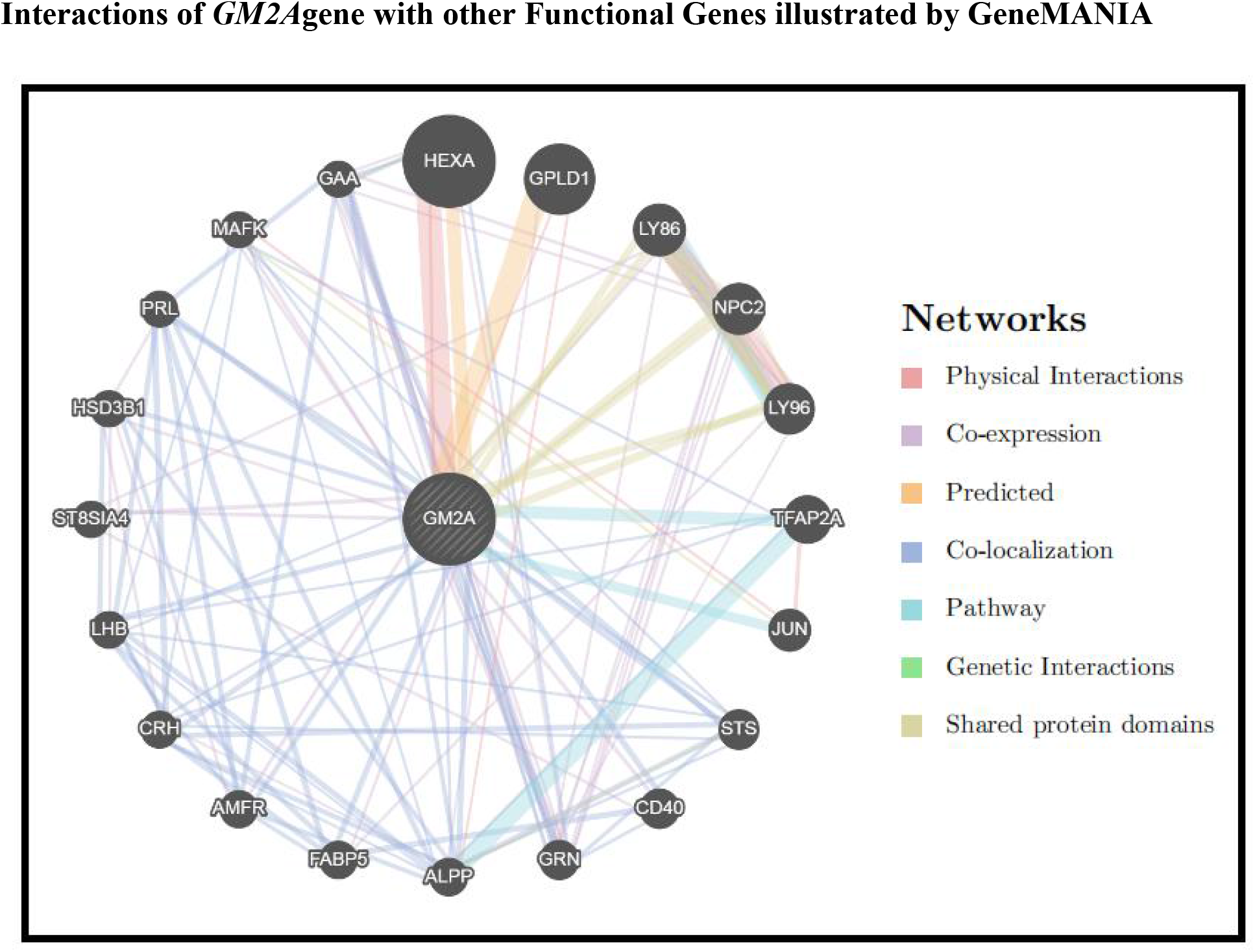
*GM2A* Gene Interactions and network predicted by GeneMania.

**Table (5):** The *GM2A* gene functions and its appearance in network and genome

| Function | FDR | Genes in network | Genes in genome |
| --- | --- | --- | --- |
| glycolipid metabolic process | 0.000729126 | 5 | 91 |
| membrane lipid metabolic process | 0.002107403 | 5 | 140 |
| glycosphingolipid metabolic process | 0.002107403 | 4 | 55 |
| sphingolipid metabolic process | 0.018998673 | 4 | 102 |
| regulation of hormone levels | 0.566018537 | 4 | 269 |
| vacuolar part | 0.566018537 | 4 | 262 |

**Table (6):** The genes co-expressed, co localized, Physical Interacted and share a domain with IL2RG

| Gene 1 | Gene 2 | Weight | Network group | Network |
| --- | --- | --- | --- | --- |
| *GAA* | *NPC2* | 0.013373704 | Co-expression | Ramaswamy-Golub-2001 |
| *GAA* | *GRN* | 0.009285928 | Co-expression | Ramaswamy-Golub-2001 |
| *NPC2* | *LY86* | 0.006769352 | Co-expression | Wang-Maris-2006 |
| *LY96* | *LY86* | 0.006558455 | Co-expression | Wang-Maris-2006 |
| *LY96* | *NPC2* | 0.004556628 | Co-expression | Wang-Maris-2006 |
| *GRN* | *NPC2* | 0.006593361 | Co-expression | Wang-Maris-2006 |
| *MAFK* | *GM2A* | 0.012908783 | Co-expression | Wang-Maris-2006 |
| *LY96* | *LY86* | 0.009755008 | Co-expression | Mallon-McKay-2013 |
| *HSD3B1* | *GM2A* | 0.012744123 | Co-expression | Mallon-McKay-2013 |
| *HSD3B1* | *CRH* | 0.011914967 | Co-expression | Mallon-McKay-2013 |
| *HSD3B1* | *LHB* | 0.011909817 | Co-expression | Mallon-McKay-2013 |
| *ST8SIA4* | *GM2A* | 0.02893034 | Co-expression | Bild-Nevins-2006 B |
| *MAFK* | *GM2A* | 0.021602744 | Co-expression | Bild-Nevins-2006 B |
| *GRN* | *GM2A* | 0.020527184 | Co-expression | Burington-Shaughnessy-2008 |
| *AMFR* | *GM2A* | 0.020826323 | Co-expression | Burington-Shaughnessy-2008 |
| *ST8SIA4* | *CD40* | 0.01166518 | Co-expression | Burington-Shaughnessy-2008 |
| *GAA* | *GRN* | 0.021335509 | Co-expression | Burington-Shaughnessy-2008 |
| *CD40* | *GM2A* | 0.026018724 | Co-expression | Dobbin-Giordano-2005 |
| *ST8SIA4* | *GM2A* | 0.02396974 | Co-expression | Dobbin-Giordano-2005 |
| *ST8SIA4* | *LY86* | 0.01011246 | Co-expression | Dobbin-Giordano-2005 |
| *LY86* | *GM2A* | 0.014090131 | Co-expression | Bahr-Bowler-2013 |
| *GRN* | *NPC2* | 0.003417451 | Co-expression | Bahr-Bowler-2013 |
| *GAA* | *NPC2* | 0.008345172 | Co-expression | Bahr-Bowler-2013 |
| *GAA* | *GRN* | 0.007346965 | Co-expression | Bahr-Bowler-2013 |
| *LY96* | *LY86* | 0.006099948 | Co-expression | Innocenti-Brown-2011 |
| *GRN* | *LY86* | 0.006802562 | Co-expression | Innocenti-Brown-2011 |
| *GRN* | *NPC2* | 0.01666259 | Co-expression | Innocenti-Brown-2011 |
| *FABP5* | *LY96* | 0.011711845 | Co-expression | Innocenti-Brown-2011 |
| *LY96* | *NPC2* | 0.006187806 | Co-expression | Rieger-Chu-2004 |
| *GRN* | *GM2A* | 0.010561576 | Co-expression | Rieger-Chu-2004 |
| *GRN* | *NPC2* | 0.005401576 | Co-expression | Rieger-Chu-2004 |
| *GRN* | *LY96* | 0.003754711 | Co-expression | Rieger-Chu-2004 |
| *FABP5* | *GM2A* | 0.014316405 | Co-expression | Roth-Zlotnik-2006 |
| *GAA* | *GM2A* | 0.010570861 | Co-expression | Perou-Botstein-2000 |
| *ALPP* | *GPLD1* | 0.003473165 | Co-expression | Smirnov-Cheung-2009 |
| *CRH* | *ALPP* | 0.004937839 | Co-expression | Wang-Cheung-2015 |
| *GRN* | *HEXA* | 0.01568475 | Co-expression | Chen-Brown-2002 |
| *GRN* | *NPC2* | 0.016359169 | Co-expression | Chen-Brown-2002 |
| *HSD3B1* | *AMFR* | 0.018177362 | Co-expression | Chen-Brown-2002 |
| *GAA* | *HEXA* | 0.01394323 | Co-expression | Chen-Brown-2002 |
| *GAA* | *GRN* | 0.012652976 | Co-expression | Chen-Brown-2002 |
| *PRL* | *HSD3B1* | 0.019465497 | Co-expression | Wu-Garvey-2007 |
| *STS* | *GM2A* | 0.030206537 | Co-localization | Schadt-Shoemaker-2004 |
| *CD40* | *GM2A* | 0.028133877 | Co-localization | Schadt-Shoemaker-2004 |
| *GRN* | *GM2A* | 0.014236832 | Co-localization | Schadt-Shoemaker-2004 |
| *GRN* | *HEXA* | 0.009402739 | Co-localization | Schadt-Shoemaker-2004 |
| *GRN* | *STS* | 0.013237921 | Co-localization | Schadt-Shoemaker-2004 |
| *GRN* | *CD40* | 0.008915158 | Co-localization | Schadt-Shoemaker-2004 |
| *ALPP* | *GM2A* | 0.025433496 | Co-localization | Schadt-Shoemaker-2004 |
| *ALPP* | *STS* | 0.025644204 | Co-localization | Schadt-Shoemaker-2004 |
| *FABP5* | *GM2A* | 0.02687657 | Co-localization | Schadt-Shoemaker-2004 |
| *FABP5* | *CD40* | 0.027478961 | Co-localization | Schadt-Shoemaker-2004 |
| *AMFR* | *GM2A* | 0.021205518 | Co-localization | Schadt-Shoemaker-2004 |
| *AMFR* | *ALPP* | 0.017865056 | Co-localization | Schadt-Shoemaker-2004 |
| *AMFR* | *FABP5* | 0.013939995 | Co-localization | Schadt-Shoemaker-2004 |
| *CRH* | *GM2A* | 0.0229119 | Co-localization | Schadt-Shoemaker-2004 |
| *CRH* | *STS* | 0.024494056 | Co-localization | Schadt-Shoemaker-2004 |
| *CRH* | *ALPP* | 0.020280294 | Co-localization | Schadt-Shoemaker-2004 |
| *CRH* | *FABP5* | 0.022545591 | Co-localization | Schadt-Shoemaker-2004 |
| *LHB* | *GM2A* | 0.021348838 | Co-localization | Schadt-Shoemaker-2004 |
| *LHB* | *ALPP* | 0.025315747 | Co-localization | Schadt-Shoemaker-2004 |
| *LHB* | *FABP5* | 0.022165282 | Co-localization | Schadt-Shoemaker-2004 |
| *LHB* | *AMFR* | 0.013310601 | Co-localization | Schadt-Shoemaker-2004 |
| *LHB* | *CRH* | 0.023500843 | Co-localization | Schadt-Shoemaker-2004 |
| *HSD3B1* | *GM2A* | 0.01956358 | Co-localization | Schadt-Shoemaker-2004 |
| *HSD3B1* | *ALPP* | 0.018070487 | Co-localization | Schadt-Shoemaker-2004 |
| *HSD3B1* | *AMFR* | 0.017620761 | Co-localization | Schadt-Shoemaker-2004 |
| *HSD3B1* | *LHB* | 0.021880075 | Co-localization | Schadt-Shoemaker-2004 |
| *PRL* | *GM2A* | 0.026748382 | Co-localization | Schadt-Shoemaker-2004 |
| *PRL* | *STS* | 0.024578847 | Co-localization | Schadt-Shoemaker-2004 |
| *PRL* | *ALPP* | 0.023220547 | Co-localization | Schadt-Shoemaker-2004 |
| *PRL* | *FABP5* | 0.027938604 | Co-localization | Schadt-Shoemaker-2004 |
| *PRL* | *AMFR* | 0.024674127 | Co-localization | Schadt-Shoemaker-2004 |
| *PRL* | *CRH* | 0.031325996 | Co-localization | Schadt-Shoemaker-2004 |
| *PRL* | *LHB* | 0.024771485 | Co-localization | Schadt-Shoemaker-2004 |
| *GAA* | *GM2A* | 0.021490874 | Co-localization | Schadt-Shoemaker-2004 |
| *GAA* | *HEXA* | 0.015766772 | Co-localization | Schadt-Shoemaker-2004 |
| *GAA* | *GRN* | 0.011875112 | Co-localization | Schadt-Shoemaker-2004 |
| *GAA* | *ALPP* | 0.016947022 | Co-localization | Schadt-Shoemaker-2004 |
| *GAA* | *AMFR* | 0.021210423 | Co-localization | Schadt-Shoemaker-2004 |
| *GAA* | *PRL* | 0.022300243 | Co-localization | Schadt-Shoemaker-2004 |
| *LY96* | *LY86* | 0.006775627 | Co-localization | Johnson-Shoemaker-2003 |
| *STS* | *GM2A* | 0.015290882 | Co-localization | Johnson-Shoemaker-2003 |
| *GRN* | *GM2A* | 0.017155321 | Co-localization | Johnson-Shoemaker-2003 |
| *ALPP* | *GM2A* | 0.01002972 | Co-localization | Johnson-Shoemaker-2003 |
| *ALPP* | *TFAP2A* | 0.009357058 | Co-localization | Johnson-Shoemaker-2003 |
| *ALPP* | *STS* | 0.008613536 | Co-localization | Johnson-Shoemaker-2003 |
| *CRH* | *GM2A* | 0.008725192 | Co-localization | Johnson-Shoemaker-2003 |
| *CRH* | *TFAP2A* | 0.008326863 | Co-localization | Johnson-Shoemaker-2003 |
| *CRH* | *STS* | 0.007740852 | Co-localization | Johnson-Shoemaker-2003 |
| *CRH* | *ALPP* | 0.005179046 | Co-localization | Johnson-Shoemaker-2003 |
| *LHB* | *GM2A* | 0.008484612 | Co-localization | Johnson-Shoemaker-2003 |
| *LHB* | *TFAP2A* | 0.008305473 | Co-localization | Johnson-Shoemaker-2003 |
| *LHB* | *STS* | 0.00752627 | Co-localization | Johnson-Shoemaker-2003 |
| *LHB* | *ALPP* | 0.005130447 | Co-localization | Johnson-Shoemaker-2003 |
| *LHB* | *CRH* | 0.004576681 | Co-localization | Johnson-Shoemaker-2003 |
| *MAFK* | *GM2A* | 0.010018589 | Co-localization | Johnson-Shoemaker-2003 |
| *MAFK* | *TFAP2A* | 0.009307076 | Co-localization | Johnson-Shoemaker-2003 |
| *MAFK* | *STS* | 0.007400996 | Co-localization | Johnson-Shoemaker-2003 |
| *MAFK* | *ALPP* | 0.005228889 | Co-localization | Johnson-Shoemaker-2003 |
| *MAFK* | *CRH* | 0.004663254 | Co-localization | Johnson-Shoemaker-2003 |
| *MAFK* | *LHB* | 0.004635511 | Co-localization | Johnson-Shoemaker-2003 |
| *GAA* | *HEXA* | 0.014947882 | Co-localization | Johnson-Shoemaker-2003 |
| *GAA* | *GRN* | 0.016629409 | Co-localization | Johnson-Shoemaker-2003 |
| *LY96* | *LY86* | 0.1184115 | Pathway | Wu-Stein-2010 |
| *TFAP2A* | *GM2A* | 0.082734935 | Pathway | Wu-Stein-2010 |
| *JUN* | *GM2A* | 0.061802242 | Pathway | Wu-Stein-2010 |
| *ALPP* | *TFAP2A* | 0.12819351 | Pathway | Wu-Stein-2010 |
| *LY96* | *LY86* | 0.1488301 | Pathway | REACTOME |
| *LY96* | *LY86* | 0.70710677 | Physical Interactions | IREF-DIP |
| *JUN* | *TFAP2A* | 0.069114916 | Physical Interactions | Li-Chen-2015 |
| *HEXA* | *GM2A* | 1 | Physical Interactions | BIOGRID-SMALL-SCALE-STUDIES |
| *MAFK* | *JUN* | 0.03714322 | Physical Interactions | IREF-BIND |
| *HEXA* | *GM2A* | 0.77715117 | Physical Interactions | IREF-HPRD |
| *GPLD1* | *GM2A* | 0.23492727 | Physical Interactions | IREF-HPRD |
| *ALPP* | *GPLD1* | 0.21245266 | Physical Interactions | IREF-HPRD |
| *HEXA* | *GM2A* | 0.41421357 | Predicted | Wu-Stein-2010 |
| *GPLD1* | *GM2A* | 0.76536685 | Predicted | Wu-Stein-2010 |
| *LY86* | *GM2A* | 0.1993388 | Shared protein domains | INTERPRO |
| *NPC2* | *GM2A* | 0.1993388 | Shared protein domains | INTERPRO |
| *NPC2* | *LY86* | 0.054153882 | Shared protein domains | INTERPRO |
| *LY96* | *GM2A* | 0.19933881 | Shared protein domains | INTERPRO |
| *LY96* | *LY86* | 0.054153886 | Shared protein domains | INTERPRO |
| *LY96* | *NPC2* | 0.054153886 | Shared protein domains | INTERPRO |
| *ALPP* | *STS* | 0.018784788 | Shared protein domains | INTERPRO |
| *MAFK* | *JUN* | 0.025053639 | Shared protein domains | INTERPRO |
| *GAA* | *HEXA* | 0.009137378 | Shared protein domains | INTERPRO |
| *LY86* | *GM2A* | 0.33333334 | Shared protein domains | PFAM |
| *NPC2* | *GM2A* | 0.33333334 | Shared protein domains | PFAM |
| *NPC2* | *LY86* | 0.33333334 | Shared protein domains | PFAM |
| *LY96* | *GM2A* | 0.33333334 | Shared protein domains | PFAM |
| *LY96* | *LY86* | 0.33333334 | Shared protein domains | PFAM |
| *LY96* | *NPC2* | 0.33333334 | Shared protein domains | PFAM |

## 4. DISCUSSION

According to our analysis we found four SNPs in GM2A gene possibly regarded as highly damaging SNPs which they are: C99Y, C112F, C138S and C138R. All of them are novel SNPs except C138R which is reported in previous studies. Sheth et al (2016) and Kochumon et al (2017) studies are agrre with us that C138R found in GM2A gene and associated with AB variant of GM2 gangliosidosis (22, 45).

Regarding C99Y (rs751417546) SNP, it has differences between wild-type and mutant residue in size and hydrophobicity; according to size, mutant residue is bigger than the wild-type residue that is located on the surface of the protein so mutation of this residue can disturb interactions with other molecules or other parts of the protein. On other hand, the wild-type residue is more hydrophobic than the mutant residue and the mutation might cause loss of hydrophobic interactions with other molecules on the surface of the protein. In case of C112F (rs773743799) SNP, the wild type residue is bigger than mutant residue also it buried in the core of a domain so the differences between the wild and mutant residue might disturb the core structure of this domain. Concerning C138S (rs1174735558) SNP, the wild-type residue is more hydrophobic than the mutant residue. The mutation might cause loss of hydrophobic interactions with other molecules on the surface of the protein.The mutation is located in a region with known splice variants. A mutation to “Arginine” was found at this position as C138R (rs137852797) which has dissimilarities between wild-type and mutant amino acids in size, charge and hydrophobicity. As the wild-type residue charge was NEUTRAL, the mutant residue charge is POSITIVE. The mutation introduces a charge at this position; this can cause repulsion between the mutant residue and neighboring residues. And According to size, the mutant residue is bigger than the wild-type residue. The residue is located on the surface of the protein; mutation of this residue can disturb interactions with other molecules or other parts of the protein. On other hand, the wild-type residue is more hydrophobic than the mutant residue. The mutation might cause loss of hydrophobic interactions with other molecules on the surface of the protein.

The wild-type residues of the four SNPs are annotated to be involved in a cysteine bridge, which is important for stability of the protein, only cysteines can make this type of bonds. The mutation causes loss of this interaction and will have a severe effect on the 3D-structure of the protein. Together with loss of the cysteine bond, the differences between the old and new residue can cause destabilization of the structure. Also the wild-type residues are extremely conserved, based on conservation scores these mutations are probably damaging to the protein.

46 SNPs out of 676 SNPs were predicted to affect miRNAs binding sites on 3’UTR leading to abnormal expression of the resulting protein. 112 alleles disrupt a conserved miRNAs binding sites while 57 derived alleles creating a new binding site of miRNAs.

*GM2A* gene function and activities illustrated by GENEMANI as following: glycolipid metabolic process, glycosphingolipid metabolic process, membrane lipid metabolic process, sphingolipid metabolic process, vacuolar part.

As the great role of using bioinformatics tools, laboratory analysis accompanied with in vivo analysis remains highly recommended confirming our findings. These findings could be used in a helpful way to improve the diagnosis and screening of AB variant of GM2 gangliosidosis.

## 5. CONCLUSION

Through using different bioinformatics tools, four SNPs were found to be highly damaging SNPs that affect function, structure and stability of *GM2A* protein, which they are: C99Y, C112F, C138S and C138R, three of them are novel SNPs (C99Y, C112F and C138S). Also 46 SNPs were predicted to affect miRNAs binding sites on 3’UTR leading to abnormal expression of the resulting protein. 112 alleles disrupt a conserved miRNAs binding sites while 57 derived alleles creating a new binding site of miRNAs. These SNPs should be considered as important candidates in causing AB variant of GM2 gangliosidosis and may help in diagnosis and genetic screening of the disease.

## 6. ACKNOWLEDGMENT

The authors desire to acknowledge the exciting collaboration of Africa City of Technology – Sudan.

## 7. DATA AVAILABILITY

All relevant data used to support the findings of this study are included within the manuscript and supplementary information files.

